# Site-specific sulfations regulate the physicochemical properties of papillomavirus-heparan sulfate interactions for entry

**DOI:** 10.1101/2024.02.08.579286

**Authors:** Fouzia Bano, Laura Soria-Martinez, Dominik van Bodegraven, Konrad Thorsteinsson, Anna M. Brown, Ines Fels, Nicole L. Snyder, Marta Bally, Mario Schelhaas

## Abstract

Certain human papillomaviruses (HPVs) are etiological agents for several anogenital and oropharyngeal cancers. During initial infection, HPV16, the most prevalent cancer-causing type, specifically interacts with heparan sulfates (HS), not only enabling initial cell attachment but also triggering a crucial conformational change in viral capsids termed structural activation. It is unknown, whether such HS-HPV16 interactions depend on HS sulfation patterns. Thus, we probed potential roles of HS sulfations using cell-based functional and physicochemical assays, including single molecule force spectroscopy. Our results demonstrate that N-sulfation of HS is crucial for virus binding and structural activation by providing high affinity sites, and that additional 6O-sulfation is required to mechanically stabilize the interaction, whereas 2O-sulfation and 3O-sulfation are mostly dispensable. Together, our findings identify the contribution of HS sulfation patterns to HPV16 binding and structural activation and reveal how distinct sulfation groups of HS synergize to facilitate HPV16 entry, which, in turn, likely influences the tropism of HPVs.

**Teaser:** Distinct heparan sulfations facilitate HPV16 binding and structural activation through their physico-chemical properties.

## Introduction

Heparan sulfate proteoglycans (HSPGs) are plasma membrane and extracellular matrix (ECM) constituents of animal cells. They consist of a protein core and long, polydisperse chains of carbohydrates, the glycosaminoglycan (GAG) HS, which varies in length and pattern of sulfation of the disaccharide core unit (*1*). Sulfation patterning is established by non-template synthesis starting with production of unsulfated, N-acetylated (NA) domains mainly composed of glucuronic acid (GlcA) and N-acetylglucosamine (GlcNAc) repeats. A series of enzymatic modifications lead to N-deacetylation and N-sulfation of GlcNAc, C5 epimerization of GlcA to iduronic acid (IdoA), and addition of O-sulfate groups at three different positions, i.e., 2O, 3O, and 6O (Fig. 1A). These synthetic steps are controlled by enzyme isoforms and abundance resulting in glycan chains that are highly heterogeneous in terms of their position and level of sulfation. Further complexity arises on a macromolecular level by separation of highly sulfated negatively charged domains (NS) and unsulfated (NA) domains.

**Fig. 1.**
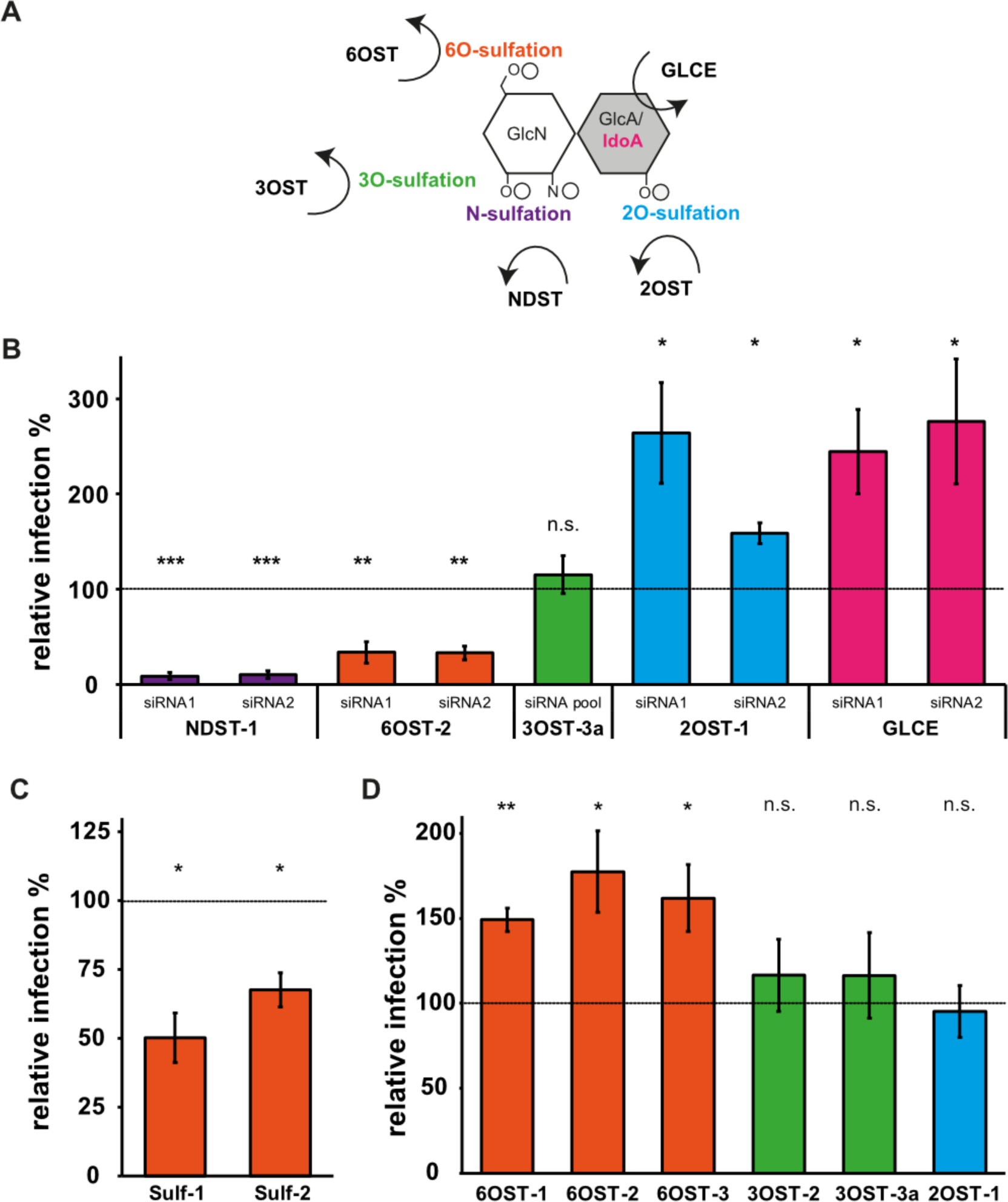
Changes in HS modifications by RNAi or overexpression of modifying enzymes affect HPV16 infectivity. (**A**) Representation of a HS disaccharide with its modifications and the modifying enzymes, with OST=O-sulfotransferase, NDST=N-deacetylsulfotransferase, GLCE=glucuronic acid epimerase, GlcN=N-acetylglucosamine, GlcA=Glucuronic acid, IdoA=Iduronic acid, circles represent sulfations. (**B**) HeLa cells were transfected with siRNAs against NDST1 and 2 (10 nM), 6OST-2 (10 nM), 3OST-3a (15 nM, pool), 2OST-1 (10 nM), and GLCE (10 nM). Two days post transfection (p.t.), cells were infected with HPV16 PsV and fixed 48h post infection (p.i.). Shown are average infection levels relative to control siRNA (10 nM, dotted line, 100%) ±standard deviation (SD). (**C**) HeLa cells were transfected with expression constructs of myc-tagged sulfatase 1 and 2. Cells were infected 24h p.t., and fixed 48h p.i.. Subsequently, cells were stained against myc, and analyzed by flow cytometry, where transfected cells were gated and infection was scored as compared to the transfected control (dotted line, 100%). Displayed are averages of three independent experiments ± S.D. (**D**) Hela cells were transfected with eGFP-tagged constructs of 6OST-1, 2, 3, 3OST-2, −3a, and 2OST-1. Cells were infected 24h p.t. and fixed 48h p.i.. Infection of transfected cells was scored by microscopy and is shown as the average of the relative infection (normalized to an eGFP transfection control) of three independent experiments. For all quantifications, a two-tailed student’s t-test was performed with p < 0.05 (*), 0.01 (**), 0.005 (***), 0.0001 (****) or non-significant (n.s.).

HSPGs are involved in a plethora of physiological processes, where biological functions are exerted through sequestration and clustering of a wide range of HS-binding proteins. This property is also exploited by pathogens; accordingly, HS interactions play a prominent role in infectious diseases (*2, 3*). The interaction of viruses with cellular glycans displays clear specificities for certain type(s) of sugars (*4*). GAGs have often been viewed to serve as rather unspecific charge-based interaction partners. However, specifically sulfated GAGs may allow virus targeting to permissive host cells via recognition of well-defined tissue-specific sulfation patterns (*5*). For example, the function of HS as an entry receptor for Herpes Simplex Virus type 1 (HSV-1) is strictly determined by the presence of 3O-sulfation (*6*). Yet, for many viruses engaging HSPGs, the role of sulfation patterning remains obscure.

Human papillomaviruses (HPVs) engage HSPGs during initial infection (*7*). HPVs are a large family of small non-enveloped DNA viruses with considerable impact on global health. Among more than 100 types, high-risk HPV16 and 18 are responsible for more than 70% of cervical cancers. Further malignancies caused by HPVs such as cancers of the head and neck display increasing incidence rates(*8*). HPVs type-specifically infect keratinocytes of either mucosa or skin. Initial infection occurs in the basal layer of these epithelia, where cells are actively dividing (*9*). Viral replication is tightly linked to epithelial differentiation, which orchestrates the expression of viral genes until newly formed viral particles are assembled in the outermost, terminally differentiated cells and released by natural shedding of dead cells (*8*).

HPV virions consist of an icosahedral (T=7) capsid formed by 72 homo-pentamers of the major capsid protein L1 and up to 72 molecules of the minor capsid protein L2, which is located mostly within a cavity of the pentameric L1 capsomers, and a double-stranded, chromatinized DNA genome of about 8kB (*10, 11*). HPV16 is one of the best studied types and often serves as a paradigm for other papillomaviruses. Due to the complex viral life cycle, growth of native viral particles is challenging. For this reason, a surrogate system termed pseudoviruses (PsVs) has been developed. PsVs consist of an L1/L2 capsid harboring a reporter plasmid (e.g. expressing GFP), which indicates successful initial infection (*12*).

Besides relying on cell-surface HS for initial interaction with host cells (*7, 13, 14*), HPVs may also bind to laminin-332 as a transient receptor in the ECM (*15*). Several HS binding sites have been identified on the pentameric HPV16 L1 capsomer (*16*). Initial attachment occurs through binding sites at the top of the pentamer, while further binding sites within the canyon between pentamers are hypothesized to be sequentially engaged (*17*). Attachment to HS triggers a conformational change, termed structural activation, resulting in the exposure of particular L1 epitopes (*13, 17*). This change is induced by highly sulfated HS, but not by other similarly sulfated glycans capable of engaging HPV16, such as chondroitin sulfate (*13*). Subsequently, further structural changes involving proteolytic cleavage of both L1 and L2 eventually lead to reduced affinity to HS and engagement of an elusive secondary receptor (complex) (*18–22*).

To date, the role of different sulfation patterns in determining the physico-chemical characteristics of HPV16-HS interactions remains understudied. Moreover, molecular details of structural activation are mostly unclear. It remains to be elucidated how activation is induced by HS-L1 interaction, and how it is manifested in structural and functional terms for the viral capsid. However, it is plausible that the underlying reason why only HS among similarly sulfated polysaccharides is able to structurally activate the virus, lies within the particular HS structure.

In this work, we investigated the interaction of HPV16 with HS on the level of distinct HS sulfation sites. Using biochemical and virological assays, our work identifies the structural constrictions within HS moieties for interaction with the capsids of HPV16, demonstrating that specific sulfations are required for HPV16 binding and structural activation, and thus infection. Furthermore, we delineate distinct roles of individual sulfation types by directly quantifying the binding kinetics and strengths for HPV16 engagement using dynamic force spectroscopy.

## Results

### RNAi of cellular sulfotransferases suggests the importance of distinct sulfations for HPV16 infection

To initially assess whether HPV16 requires specific sulfations for HS engagement, distinct, highly expressed sulfotransferase isoforms (fig. S1A) were successfully depleted individually by RNAi in HeLa cells (fig. S1B), after which cells were infected. RNAi of N-deacetylase sulfotransferase 1 (NDST-1), the majorly expressed NDST isoform, strongly impaired HPV16 infection by about 90% (Fig. 1B). Since NDSTs exchange acetyl to sulfate groups in GlcNAc, this indicated the importance of N-sulfation. 6OST-2 depletion reduced infection by about 70% (Fig. 1B), indicating that 6O-sulfation is also important in infection. Interestingly, reduced levels of 2OST-1, responsible for 2O-sulfation, and glucuronic acid epimerase (GLCE), mediating isomerization from glucuronic to iduronic acid, increased infection by up to 2.5-fold suggesting that these modifications could be unfavorable (Fig. 1B). 3O-sulfation is a rare glucosamine modification of HS (*23*). Thus, expression of 3OSTs was expectedly low: only 3OST-2 and −3a were significantly expressed (fig. S1A). Depletion of 3OST-2 caused major cell toxicity. 3OST-3a could be depleted using a pool of three siRNAs (Fig. 1B, fig. S1B). This led to functional reduction of 3O-sulfation as validated by reduced HSV-1 infection (fig. S1C, (*6*)), but not to reduced HPV16 infectivity (Fig. 1B), indicating that 3O-sulfation is not a major determinant for HPV16 infection. In corroboration, 3O-sulfation-deficient chinese hamster ovary (CHO) cells, were readily infected by HPV16 independent of 3OST expression (fig. S1D), while HSV-1 infection was only successful if 3OST-3b was ectopically expressed (fig. S1D). This confirmed that 3O-sulfation is not strictly required for HPV16 infection. In summary, the data indicated that N- and 6O-sulfation are important for infection, whereas further sulfations or structural features are not crucial or even unfavorable for infection. It is important to note, however, that the degree of specific sulfations is not directly correlated with the presence of the enzymes but may also depend on the presence or absence of other sulfations, e.g. O-sulfation typically does not occur in the absence of N-sulfation, or 2O-sulfation is much more efficient after epimerization of GlcA to IdoA(*1*).

To verify the notion that specific HS sulfations facilitate or perhaps restrict HPV16 infection, we overexpressed sulfatases or sulfotransferases prior to infection. Sulfatase (Sulf) 1 or 2 (Fig. 1C) specifically remove 6O-sulfates extracellularly thus reducing 6O-sulfation levels of HS. Sulf-1 and −2 overexpression decreased infection by around 50% and 35%, respectively (Fig. 1C). In turn, overexpression of the different 6OST isoforms resulted in 50% to 70% increased infection (Fig. 1D), whereas overexpression of 2OST-1 or 3OST-3a failed to affect HPV16 infection.

Collectively, our results from RNAi and overexpression experiments indicated that N- and 6O-sulfation in HS are important for HPV16 infection, whereas 2O- and 3O-sulfation are dispensable.

### Role of individual sulfations in HPV16 capsid binding to heparin

To more specifically examine the importance of individual sulfations within HS, competition assays with chemically desulfated and parental (unmodified) heparins (methods and Table S1 for details) were employed. Heparin is a secreted, almost fully sulfated version of HS, and is commonly used to represent the NS domains of HS. Here, HPV16 was preincubated with parental and selectively desulfated heparin polysaccharides of the same batch and therefore the same dispersity with only specific sulfations being absent or present. We focused on polysaccharides, as small oligosaccharides engaging individual binding site only exhibit limited affinity, suggesting multivalent binding as key for efficient engagement (*13, 24*). Only a stable binding of the heparin in question to the capsid prior to virus addition to cells could then interfere with infection as it prevents binding to cellular HS (*13*). Of note, the importance of 3O-sulfation could not be probed in such experiments, as it cannot be specifically chemically desulfated. Expectedly, parental heparin completely prevented infection (Fig. 2A, (*13*)) and binding of viral particles to cells (Fig. 2B, (*13*)), indicating that fully sulfated heparin interacted very stably with HPV16 capsids. 2O-desulfated heparin (deS) also dose-dependently blocked infection and impaired binding (Fig. 2A-C). However, 2-log higher concentrations were needed to achieve similar blocking as parental heparin. Hence, 2O-sulfation was not strictly required for HPV16 binding to HS, although it may contribute to interaction stability. This finding also implies that increased infection upon reduced 2O-sulfation after knockdown was caused by a pleiotropic effect, perhaps by affecting cellular processes restricting infection rather than virus binding. In contrast, 6O-deS and N-deS were unable to block infection and cell-binding of HPV16 (Fig. 2A-C), indicating that these heparins could not compete with cell-surface HSPGs for binding. Thus, 6O-sulfation and N-sulfation were crucial for stable binding of heparin to HPV16 delineating the importance of individual sulfations.

**Fig. 2.**
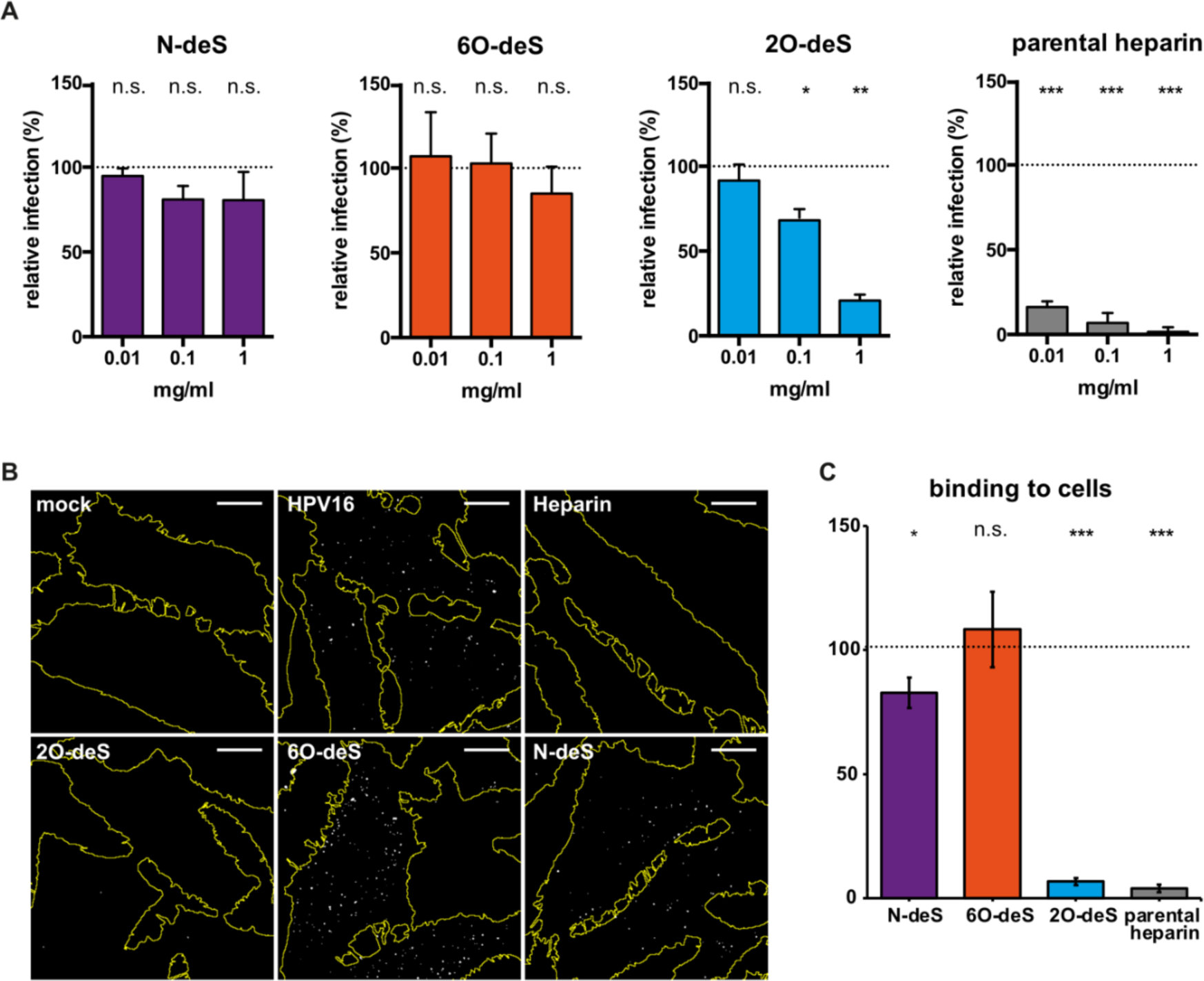
N-sulfation and 6O-sulfation are required for stable interaction with HPV16 capsids. (**A**) To test for competition of desulfated and parental heparins for binding to cells (add-on assay), HPV16 PsV were incubated with polysaccharides for 1h at RT. This inoculum was added to cells and allowed to bind for 2h at 37°C. After washing off unbound virus, infection was allowed to proceed until 48h p.i., when cells were fixed and infection was scored by automated microscopy. Shown are infection levels relative to an untreated HPV16 control (dotted line, 100%) from three independent experiments ± SD. (**B**) Desulfated/parental heparins were preincubated at a concentration of 1mg/ml with fluorescently-labeled HPV16. Two hours after addition of the inoculum to cells, cells were fixed and analyzed by confocal microscopy. Shown are maximum intensity projections of confocal stacks with cell outlines in yellow and virus particles in white. Scale bar is 10µm. (**C**) Virus binding on cells was quantified using the Fiji 3D object counter plugin. The signal of cell-associated virus is depicted normalized to an untreated HPV16 control (dotted line). depicted are averages of three independent experiments ±SD. For all quantifications, a two-tailed student’s t-test was performed with p < 0.05 (*), 0.01 (**), 0.005 (***), 0.0001 (****) or non-significant (n.s.).

### N- and 6O-sulfation facilitate structural activation of HPV16

Besides mediating attachment, the interaction of HPV16 with HS also induces a structural change in the capsid (*13*). Thus, we evaluated whether the structural constraints of HS for binding extended to structural activation, using a functional assay for structural activation. This assay relies on attaching HPV16 to laminin-332 through HS-independent binding sites, followed by seeding of HS-undersulfated cells on top (Fig. 3A). These cells are only infected, if the virus is preincubated with an HS that can elicit the conformational change leading to structural activation (*13, 25, 26*). In this assay, parental and 2O-deS heparin recovered infection of NaClO_3_-treated cells (*13*) to a similar extent, indicating that 2O-sulfation was dispensable for structural activation (Fig. 3B). In contrast, N-deS failed to structurally activate the virus (Fig. 3B), highlighting the key role of N-sulfation in structural activation. 6O-deS could still partially activate HPV16, although only to about 50% of the effect of parental heparin (Fig. 3B). Since ECM-binding of the virus was unaffected by glycan binding as expected (fig. S2, (*13*)), any impairment of infection was caused by failed structural activation. Hence, structural activation was controlled and facilitated by N- and to a lesser extent by 6O-sulfation, while 2O-sulfation was dispensable.

**Fig. 3.**
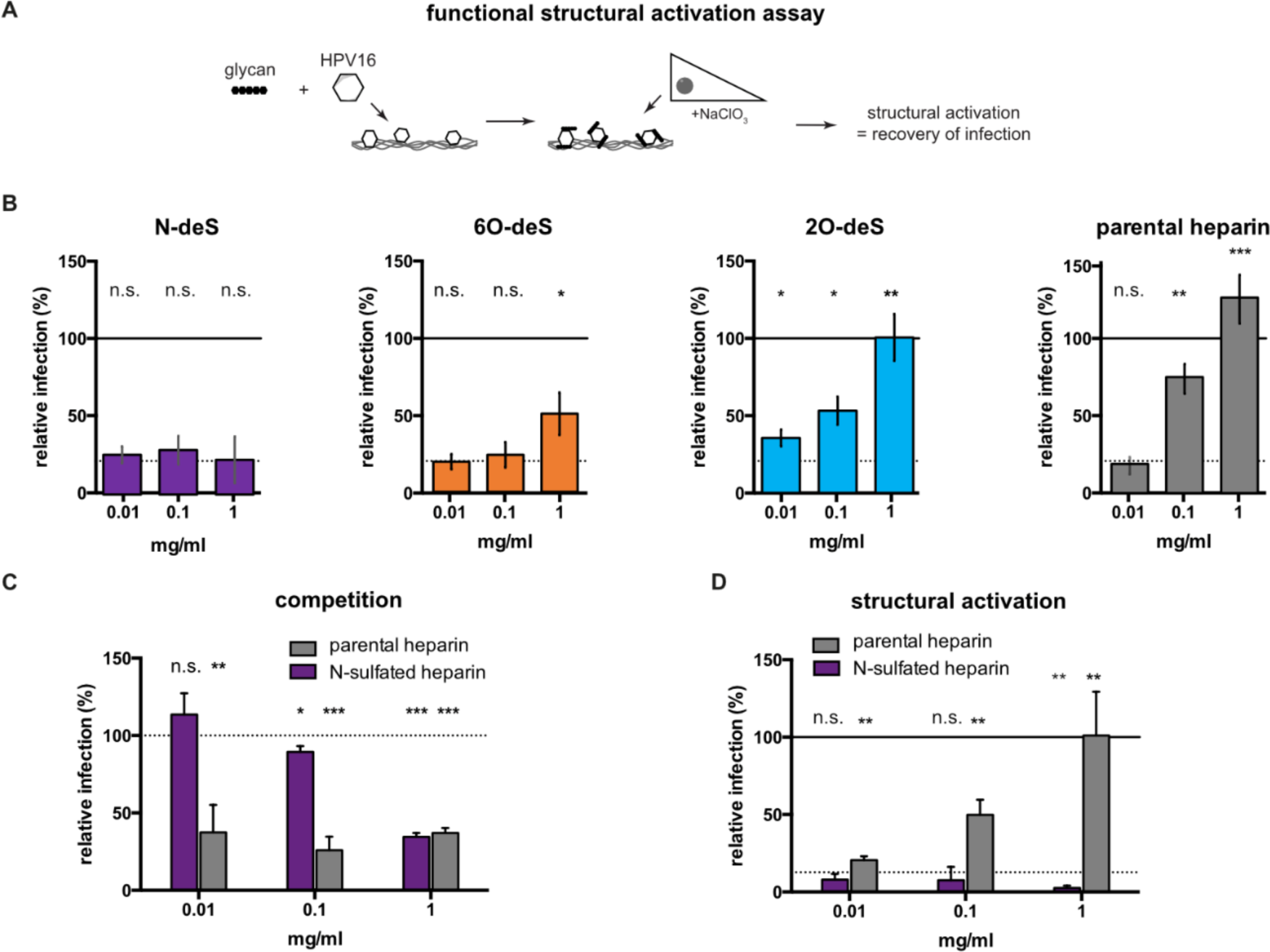
N- and 6O-sulfation contribute to structural activation of HPV16. (**A**) Schematic representation of the ‘seed-over assay’ to test for structural activation of HPV16. (**B**) HPV16 PsV were pre-incubated with desulfated and parental heparins for 1h at RT and allowed to bind to undersulfated extracellular matrix (ECM) derived from HaCaT cells for another hour. Infection of over-seeded, undersulfated HeLa cells was scored 48h p.i. as a readout for structural activation. Shown are infection levels relative to untreated, add-on infection of HPV16 (full line), whereas background infection levels of the undersulfated, NaClO_3_-treated cells in seed-over condition is indicated by a dotted line. Depicted are averages of three independent experiments ± SD. (**C**) and (**D**) Exclusively N-sulfated heparin and the parental fully sulfated heparin were used in add-on (C) and seed-over (D) assays. Infection levels are shown relative to untreated HPV16 (full line) ±SD for three independent experiments. In C, the dotted line represents background infection of NaClO_3_ treated cells as in B). For all quantifications, a two-tailed student’s t-test was performed with p < 0.05 (*), 0.01 (**), 0.005 (***), 0.0001 (****) or non-significant (n.s.).

### N-sulfation alone is insufficient for structural activation of HPV16

Since N-sulfation was essential for binding and structural activation of HPV16 and 6O-sulfation appeared to play a supportive role, we wondered whether N-sulfation alone could be sufficient. To test this, exclusively N-sulfated heparin was generated by chemically desulfating heparin completely and subsequently chemically re-sulfating only N-residues (see material and methods). While this derivative blocked HPV16 infection, it required a 2-log increase in concentration compared to parental heparin (Fig. 3C) indicating a role of further sulfations for stable engagement. Moreover, N-sulfated heparin failed to structurally activate HPV16 (Fig. 3D). Thus, N-sulfation alone was insufficient, and the presence of additional sulfations (most likely 6O-sulfation) was required for structural activation.

### N-sulfation is essential for high affinity interaction of HPV16 with heparin

To understand how individual sulfation groups contribute to HPV16 binding and structural activation, we next explored the physical nature of HPV16-heparin interactions by atomic force microscopy (AFM)-based single molecule force spectroscopy (SMFS). Towards this, we implemented and characterized a glycocalyx mimic platform consisting of heparin chains coupled to the surface via a biotin-tag at their reducing end (*27, 28*) (Fig. 4A). Desulfation did not significantly affect the physical properties of the films or the heparin chain length (see Table S1 and S2, fig. S3, and supplementary text S4 for characterization).

**Fig. 4:**
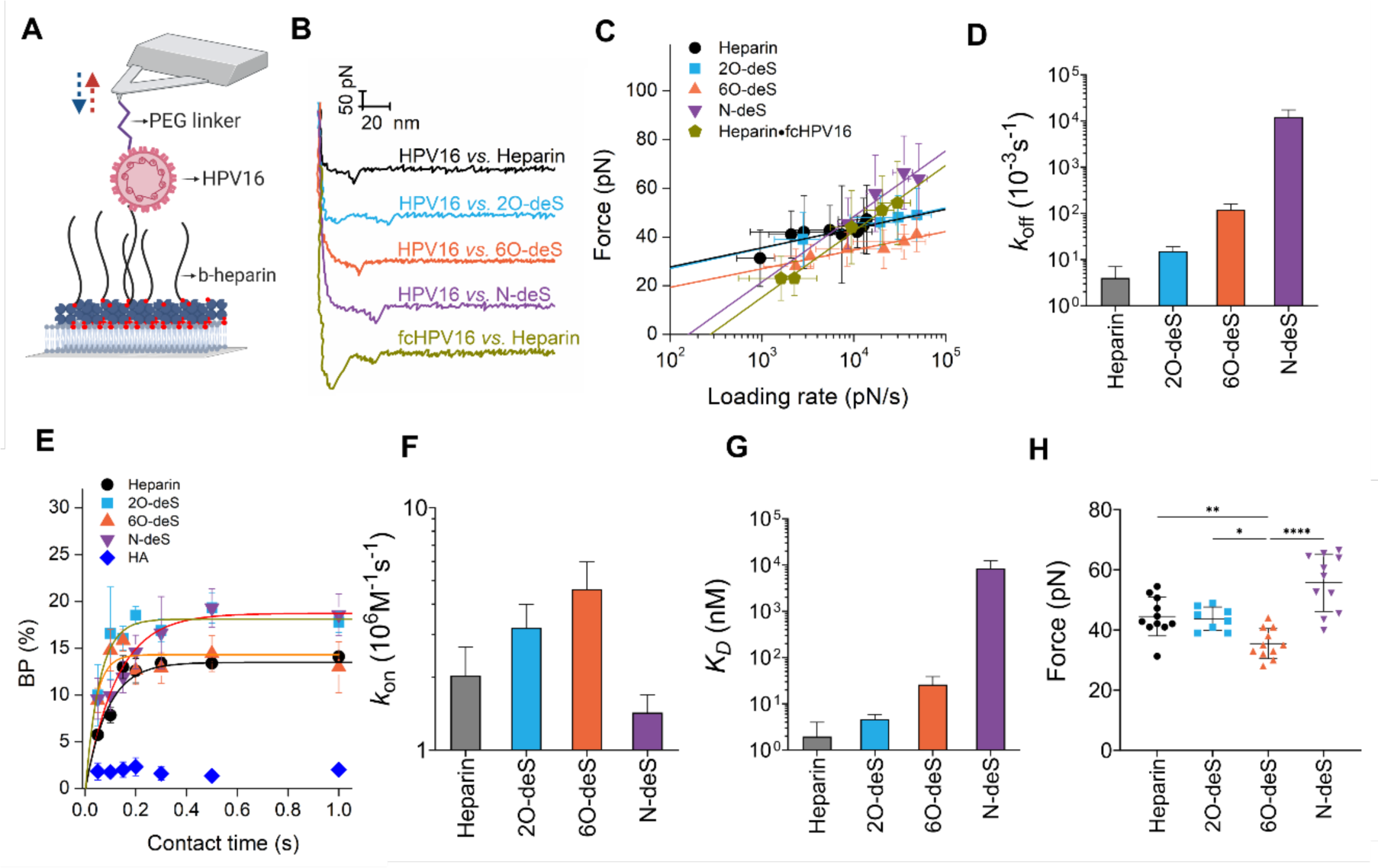
Quantifying HPV16-heparin bonds by AFM-based SMFS. (**A**) Schematic illustration of the experimental setup of AFM-based SMFS. A single HPV16 virion is attached to an AFM probe via a PEG linker and brought into contact with a heparin-presenting surface for the repeated recording of extend and retract curves. (**B**) Representative retract curves recorded at 1 µm/s with a maximum applied load of 600 pN for specific unbinding events between HPV16, or fcHPV16 and indicated heparin surfaces. (**C**) Dynamic force spectra covering a range between 1×10^3^ and 5×10^5^ pN/s for HPV16 interactions with four heparin surfaces, and fcHPV16 interaction with heparin, as indicated. Data is representative from at least 2 independently prepared AFM probes and heparin surfaces. Solid symbols represent mean rupture force obtained from Suppl. Fig. 7 and error bars represent standard deviations. The solid lines are best fits to the Bell-Evans model. (**D**) *k*_off_values extracted from the fits. (**E**) Plots of BP as a function of contact time are shown for the indicated HPV16-heparin interactions. Solid lines are the best fits to an exponential function (Eq. 2) to extract the interaction time (τ) needed to reach half maximum BP. Error bars represent the standard error of the mean (SEM) from 3 independent experiments. (**F**) *k*_on_ values derived from BP plots. (**G**) *K*_D_ values derived from *k*_off_ and *k*_on_ from (D) and (F), respectively. (**H**) Box and whisker plot with 10-90 percentile of unbinding force data for 4 heparin surfaces for data points = 1204 (heparin), 971 (2O-deS), 1088 (6O-deS) and 1217 (N-deS) from 3 (heparin and N-deS) and 2 (2O-deS and 6O-deS) independent experiments. P-values were determined by one-way ANOVA test in GraphPad Prism (Brown-Frosythe and Welch ANOVA test). P<0.0001 (****), P = 0.0033 (**) and P = 0.0119 (*).

We assessed the influence of different sulfate groups on the binding affinity between HPV16 and heparin, by measuring kinetic off and on rates using AFM-based SMFS. In these experiments, an AFM tip carrying a single HPV16 virion immobilized via a polyethylene glycol (PEG) linker (Fig. 4A, fig. S4) was approached to the heparin surface, allowing the virion to interact with heparin. Subsequently, the tip was retracted until the external force pulling on the bond exceeded its mechanical stability, resulting in rupture. Retract curves from the obtained force distance (FD) curves for all tested conditions displayed the typical features of the elastic stretching of a single PEG chain followed by a rupture event (Fig. 4B). From these rupture events, force histograms for unbinding forces were derived (fig. S5A) after verifying that the interactions are taking place between a single virion and a single heparin chain, which we will refer to as single-molecule interactions in the rest of the text (see supplementary materials for details).

Having verified the specific nature of HPV16-heparin in several control experiments (supplementary materials S4), we then extracted intrinsic kinetic bond parameters (e.g., kinetic off-rates at zero force, *k*_off_) by probing the interactions at various loading rates and fitting the data with the Bell-Evans model (Fig. 4C, fig. S7, Eq. 1, see methods for details). Briefly, the Bell-Evans model proposes a single energy barrier between bound and unbound state if there is a linear dependence of the force on the logarithm loading rate, which was the case for all the interactions tested here. A comparison of the extracted kinetic parameters first indicated that HPV16 dissociates from N-deS at a much higher rate as compared to all other interactions studied here (Table S3). N-deS exhibited a *k*_off_ value of 12 ± 5 s^-1^, i.e., about 3000 times lower than parental heparin, implying unstable interactions in the absence of N-sulfates. Second, this comparison showed that HPV16 unbound from 6O-deS with a *k*_off_ value that was ∼30 times lower than parental heparin, whereas this value was only ∼4 times lower for 2O-deS as compared to parental heparin (Fig. 4D, Table S3), suggesting that 2O sulfates are the most dispensable in ensuring stable interactions. This is in line with our cell-based observations. Finally, low-HS-affinity(*18*), so-called furin-precleaved HPV16 (fcHPV16) interaction with heparin revealed similarly low kinetic off-rate with the value of 18.7 ± 6.6 s^-1^ (Fig. 4C, Table S3) as compared to HPV16 and N-deS, implying that the interactions between fcHPV16 and heparin are unstable, as expected. Collectively, our *k*_off_ data suggests that N-sulfates contribute to the interactions between HPV16 and heparin by facilitating a longer bond lifetime, with further contributions by 6O-sulfates.

We next determined the kinetic on-rates (*k*_on_) of the HPV16-heparin bond formation by experimentally measuring the influence of contact time on the binding probability (BP) at a fixed retract velocity (Fig. 4E) (see methods for details). From *k*_on_ calculations (Fig. 4F, Table S3), we observed similar or slightly slower on-rates for the HPV16-N-deS and the HPV16-heparin interaction, whereas these rates were similar or slightly higher for the case of 2O-deS and 6O-deS heparins. This indicates that the individual types of sulfation only marginally contribute to efficient association of heparin to the capsid, but rather the sum of present sulfations dictate the rate of association.

Considering both the *k*_on_ and the *k*_off_values determined here, we conclude that, expectedly, HPV16 has the highest binding affinity (*K*_D_) to parental heparin among all heparin derivatives (Figure 4G, Table S3). Notably, the *K*_D_ value for HPV16 binding to N-deS was three orders of magnitude higher than for heparin indicating that the N-sulfates are key in ensuring a high affinity interaction, primarily by increasing the bond life-time.

### Removal of 6O-sulfation of heparin mechanically weakens the HPV16-heparin interactions

Beyond quantifying the binding affinity of HPV16-heparin interactions, AFM-based SMFS data also provides information about the mechanical strength of these interactions. Here, the mean rupture forces were significantly the lowest for 6O-deS over the whole range of loading rates probed here and fell by ∼24% compared to parental heparin (Fig. 4H). This suggested that 6O-sulfation greatly contributes to the mechanical strength of HPV16-heparin interactions. Most interestingly, a higher unbinding force in fast loading regime was observed for the interaction of HPV16 with N-deS, despite its low binding affinity, implying that there is no close correlation between affinity and mechanical strength of molecular bonds (*29, 30*).

Taken together, our biophysical analyses on heparin and its derivatives suggest that, while the presence of N-sulfations is crucial in increasing the lifetime of the interactions, 6O-sulfation predominantly contributes to the mechanical strength of HPV16-heparin interactions, thus making them much more stable.

## Discussion

This work describes for the first time how different HS sulfation types aid viral invasion through separate engagement roles, by exhibiting distinct physico-chemical properties in their interaction with viral capsids. Specific sulfations within HS were crucial for stable glycan engagement and structural activation of HPV16: N-sulfation was critical yet alone insufficient, 6O-sulfation contributed to it, while 2O-sulfation was only marginally important. Our physico-chemical investigations of the glycan-virus interactions revealed that all types of sulfates contribute to initial association of virus and glycan signified by similar *k*_on_. Thus, initial association is most likely driven by charge-based electrostatic interactions depending on the number rather than on the type of sulfate groups. Yet, N-sulfation was key in stabilizing the interaction between the virus and glycans by increasing the bond’s lifetime with resulting orders of magnitude lower *K*_D_. This resistance to detachment was further strengthened by 6O-sulfation, which greatly contributed to the mechanical stability of heparin-HPV16 interactions despite its moderate effect on *K*_D_. From this emerges a model in which early sampling of the cell surface’s negatively charged glycans leads to initial virus association, and encountering N- and 6O-sulfated HS moieties strengthens the interaction by extending bond lifetime and providing transient mechanical stabilization of the bond, which leads to structural activation (Fig. 5). Eventually, the almost irreversible binding to cells of HPV16 is likely driven by avidity, i.e., engagement of multiple HS molecules by the capsids.

**Fig. 5:**
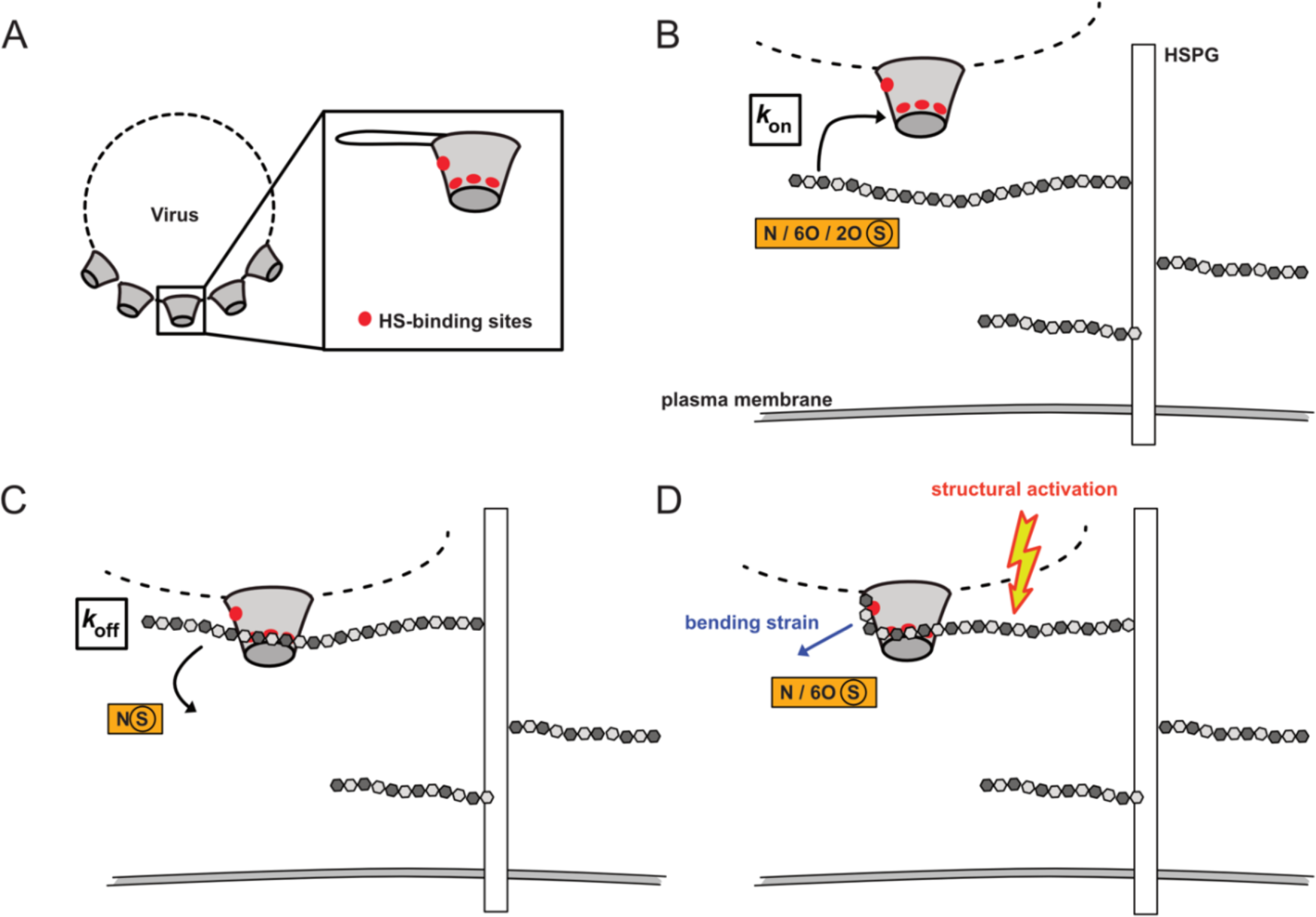
Proposed model of the contribution of HS sulfation moieties to virus binding kinetics and structural activation. (**A**) Schematic illustration of four binding sites (inset) on the HPV16 capsomer. (**B**) N-, 6O-, and 2O-sulfation moieties contribute to the association of HS. (**C**) Predominantly, N-sulfation controls unbinding. (**D**) 6O-sulfation contributes to structural activation by providing mechanical stability against HS bending strain with N-sulfation contributing low *k*_off_ rates. Please note, that the individual sulfations exert their contribution not necessarily in a sequential fashion.

Ligand affinity to defined glycans is most often used as indicator for the importance and functionality of an interaction; a direct relationship between affinity, binding and functional outcome is often assumed. However, binding strength and function do not necessarily correlate, as reflected by published work and our current results. Previous work reveals that chondroitin sulfate bind HPV16 capsids comparably to HS (*13*), can block infection (*13*), and allow (transient) binding to cells, if HS is unavailable (*31*); yet only HS can induce structural activation (*13*). Also, short heparin oligosaccharides (dp8-12) bind to capsids (*13, 16, 24*), but fail to block infection (*16*). Indeed, our force spectroscopy data evidenced several orders of magnitude higher affinities for N-vs. 6O-sulfation. Noteworthy, 6O- and 2O-sulfation contribute similarly though moderately to affinity, with *K*_D_values in the low nM range for both de-sulfated compounds. Yet, 6O-sulfation is crucial for infection, strong binding, and to a good extent for structural activation, whereas 2O-sulfation is not. This raises the question of how and why the different sulfations contribute differentially to binding and structural activation. As mentioned above, *k*_on_ rates were similar for all types of sulfation likely providing the means of initially establishing a first contact between HS and the virus based on the abundance rather than type of sulfates. N-sulfates provided low *k*_off_ rates to stabilize binding. Our data on the role of 6O-sulfates to mechanically stabilize bonds while exhibiting moderate affinity in combination with their role in infectivity, binding, and structural activation data implies that (i) the mechanical strengthening effect is of notable importance for a functional interaction, and (ii) this mechanical stability is unlikely to be determined by charge-based interactions typically contributing majorly to affinity. While this has not been observed for any virus-GAG interactions, it is not entirely unprecedented. For example, the individual bond between HA in the glycocalyx and CD44 on leukocytes, known to play a key role in cell adhesion to the endothelium, is highly mechanically stable despite its low, µM-range binding affinity (*29*), a feature supporting the significant role for CD44-HA interaction in leukocyte migration under the shear stress of blood flow.

So, why would there be a need for mechanically strong connections within the HS-virus bond in the context of HPV16 infection? Since our data suggests that 6O-sulfation is ultimately important for structural activation, a process correlated with a conformational change in the virion which allows for further enzymatic processing towards successful entry (*13, 20*), the mechanically strong connections by 6O-sulfates are perhaps key for this phenomenon. Given that the proposed HS binding sites in the capsid (*16*) line up along the top rim to face downwards into the canyon between pentamers, it is possible that a single HS molecule engages all those binding sites (Fig. 5). Indeed, our recent work strongly supports such a model for structural activation, in which the engagement of one HS molecule of the binding pockets in the capsid stabilizes one of several alternative conformations in the C-terminal arm connecting the capsomers, leads to particle softening and enlargement(*32*). However, such an engagement would necessitate a bend within the HS molecule eliciting mechanical strain on the HS-virus binding sites (Fig. 5D). Withstanding such mechanical forces is then likely conferred by 6O-sulfation’s ability to provide resistance to mechanical rupture upon glycan-virus binding. Mechanical stabilization of the bond may also be of relevance in an *in vivo* context: HPV16 infects its target cells, the basal cells of squamous epithelia, after tissue damage (*9*). Mechanical stabilization of the initial bond to HSPGs is likely to represent a particular advantage during the wound healing process, where virus particles bound to cell and/or ECM-associated HSPGs likely experience substantial mechanical force during cellular migration and/or remodeling of the glycocalyx. As a specific example, the transfer of particles from the ECM to cells by and locomotive transport along filopodia by actin retrograde transport (*33*) most certainly exerts mechanical force on the HSPG-virus bond.

Taken together, our work reveals that N- and 6O-sulfation primarily function as stabilizing and strengthening residues respectively, in the HPV16-HS interaction. This unique interplay between high-affinity interaction and resistance to tensile force provides a mechanism for GAGs-mediated virus entry and raises the question of whether our findings represent a paradigm also for other virus-GAG interactions. In future work, it would be most interesting to test whether mechanical stability of HS-virus bonds is of general importance across different virus families. Also, it would be of relevance to investigate whether different HPV types with tropisms for mucosa *vs.* skin epidermis exhibit different HS-sulfation requirements, aligned with different sulfation patterns of human tissues.

## Materials and Methods

### Cell culture

HeLa cells were from ATCC. Chinese Hamster Ovary cells (CHO)(*34*) were a kind gift from K. Grobe (WWU Münster, Germany). HaCaT cells were a kind gift of J.T. Schiller (NIH, Bethesda, USA) but originated from N. Fusenig (DKFZ, Heidelberg, Germany)(*35*). 293TT cells (*12*) were a kind gift of J.T. Schiller (NIH, Bethesda, USA). 293TTF cells (*36*) were a kind gift from R. Roden (John Hopkins University, Baltimore, MD, USA).

HeLa, HaCaT, 293TT, and 293TTF cells were cultured in DMEM (Sigma) supplemented with 10% fetal bovine serum (Capricorn) at 37°C and 5% CO_2_, whereas for CHO cells F-12 media (Sigma) was used. Cells were regularly tested for mycoplasma contamination.

### Transfections

DNA vector transfection of cells was performed one day after seeding at a confluency of 50-60% using Lipofectamine 2000 (ThermoFisher) according to the manufacturer’s instructions.

### Plasmids generation

Plasmids for sulfatases were from Addgene (#13003 and #13004). Heparan sulfate sulfotransferase genes (in pDONR223 from the ORFeome v8.1 collection (Dharmacon)) were cloned by LR reaction (ThermoFischer Scientific) into mammalian expression plasmid pDest-eGFP-N1 (Clontech) according to the manufacturer’s instructions. The construct for 3OST-3a (in pOTB7 from the Mammalian Gene Collection (MGC, Dharmacon)) was cloned by BP reaction (ThermoFischer Scientific) into entry vector pDONR221 (Invitrogen) according to the manufacturer’s instructions, and subsequently into pDest-eGFP-N1 as above.

### Virus preparation

HPV16 expressing eGFP, dsRed, or mCherry were produced using the plasmids p16sheLL(*37*) and pClneo-GFP, pClneo-dsRed, or pClneo-mCherry, respectively, in 293TT cells as previously described (*12*), whereas fc-HPV16 PsV were produced using furin-overexpressing cells (293TTF). In short, 293TT or 293TTF cells were transfected for 48 h with the corresponding plasmids, cells were harvested, and lysed. In case of fcHPV16, cell lysates were incubated for 24 h supplemented with 10 mM HEPES (pH 7.6) and 2 mM CaCl_2_. For optimal maturation, all lysates were incubated for further 24 h with 25 mM ammonium sulfate (pH 9.0). PsVs were purified on a 25%-39% linear Iodixanol (OptiPrep) gradient (Sigma-Aldrich).

Fluorophore-labeled HPV16 or furin-precleaved (fc) PsVs were prepared as described previously(*33*). In short, HPV16 PsVs were incubated for 1h with fluorophore-conjugated succinimidyl ester (Alexa Fluor 568 or 488) at a 1:8 molar ratio (L1/dye). Labeled viruses were separated from free dye by an Optiprep step gradient (5%/39%).

HSV-1 (strain 17 S17)(*38*) and HSV-1 GFP were prepared as described previously(*39*). Briefly, virus inoculum was prepared by infecting monolayers of confluent BHK cells at a multiplicity of 0.003 plaque forming units (p.f.u.) per cell at 31°C. At 24 h p.i., the inoculum was replaced by culture medium. Virus was concentrated from the culture medium at about 120 h p.i. by centrifugation for 90 min at 12 500 r.p.m.

### Infectivity assays

#### HPV16 competition assay

For competition studies between purified glycans and cell glycans, 4000 HeLa cells were seeded into 96-well optical-bottom-well plates (Greiner) prior to infection. Heparin or other polysaccharides (Table S1) were incubated at the indicated concentrations with around 8 ng HPV16 PsV for 1h at room temperature (RT) in PBS. The inoculum was added to cells and incubated at 37°C for 2h, after which the inoculum was exchanged for fresh growth media. Cells were fixed with 4% paraformaldehyde (PFA) at 48h p.i.. Cell nuclei were stained with RedDot2 and infection was scored by automated microscopy using a spinning disc microscope (Zeiss Axio Observer Z1, equipped with a Yokogawa CSU22 spinning disc module; Visitron systems GmbH, or a Nikon Ti2 eclipse equipped with a Dragonfly 600 spinning disc module, Oxford Instruments) with a 20x objective. Analysis was done by computational analysis. For each image, the total number of cells and the infected cells (GFP+) was calculated by Infection Counter, a MATLAB-based infection scoring script(*40*). The infection of each condition was normalized to untreated virus control.

#### HPV16 structural activation assay

To determine whether certain glycans structurally activate HPV16, NaClO_3_-treated HaCaT were seeded to confluency (4×10^4^) into 96-well optical-bottom-well plate (Greiner) to generate ECM with unsulfated HS. Cells were treated daily with 50mM NaClO_3_. After 48h, HaCaT cells were detached with 20 mM EDTA, and ECM was washed with PBS. The indicated GAGs were incubated at the indicated concentrations with around 15 ng HPV16 PsV for 1h at RT. Virus was then allowed to bind ECM for 1h and non-bound virus was removed by washing with PBS. 4000 HaCaT (NaClO_3_-treated or untreated) were seeded per well onto the ECM-bound HPV16. NaClO_3_-treatment was refreshed daily. At 48h p.i., cells were fixed with 4% PFA and infection scoring proceeded as for the competition assay. Infection was expressed as relative to the untreated cells infection level.

### RNAi and subsequent infection

Pre-designed small interfering RNAs (siRNAs) and the non-target control (negative control) AllstarNeg were from Qiagen. 2×10^4^ HeLa cells were reverse transfected with 10 nM siRNA using Lipofectamine RNAiMax (Life Technologies) according to the manufacturer’s instructions. For 3-OST-3b, the selected siRNAs were pooled, and cells were transfected with a final concentration of 15nM for 40h. After this, cells were trypsinized, counted and re-seeded to ensure equal number of cells upon infection. After 48h of transfection, cells were infected with HPV16 PsV or HSV-1 GFP to result in about 20% infected cells of the non-target control or cells were lysed for RNA extraction. For infection samples, cells were fixed 48h p.i. and infection was assessed as for add-on experiments and expressed as relative infection (%) to AllstarNeg.

### Infection after sulfatase and sulfotransferase overexpression

The day before transfection, 5×10^4^ HeLa cells were seeded in 12-well plates. Vectors of sulfatase 1 and 2 tagged with myc were transfected using Lipofectamine 2000 (ThermoFisher) according to the manufacturer’s instructions. Media was exchanged 6h post transfection. Cells were infected for 48h with HPV16-GFP PsVs at 24h post transfection to result in 20% infected cells for control transfected cells. Cells were trypsinized, fixed in 4% paraformaldehyde, and permeabilized using saponin-containing PBS. Cells were stained for sulfatases using an anti-myc antibody. Scoring of infection (GFP-positive cells) and transfection (myc-positive cells) was performed by flow cytometry. The number of GFP-positive cells of the myc-positive cells (infected cells) was calculated relative to the control sample.

For the transfection of HeLa and CHO cells with sulfotransferases plasmids, 5×10^3^ and 7×10^3^ cells were seeded, respectively, and transfection and infection (with HPV16-mCherry) proceeded as for sulfatases. Briefly, the mixture of plasmid and Lipofectamine was mixed with cells, which were then added to 96-well optical-bottom-well plate (Greiner). After 24h, cells were infected either with HPV16-mCherry (10 ng) and fixed 48h p.i., or with HSV-1 (1 p.f.u./cell) and fixed at 6h p.i.. RedDot2 staining was used according to the manufacturer’s instructions for staining of nuclei, whereas for HSV-1 infected cells ICP0 was stained using an ICP0 antibody and AF568-labelled secondary antibody. The plate was imaged by automatic microscopy with a spinning disc microscope (as above). Analysis was done by computational analysis. For each image, the total number of cells (redDot2+), the transfected cells (GFP+), and the number of infected cells (mCherry+) of the transfected cells (GFP+) was determined by Infection Counter, a MATLAB-based infection scoring script(*40*).

### RNA extraction and quantitative PCR

RNA extraction was performed using RNeasy Kit (Qiagen) according to supplier’s instructions. The RNA was reverse transcribed using RevertAid Reverse transcriptase (ThermoFisher) following the manufacturer’s instructions. Expression levels were measured by quantitative PCR using pre-designed primers (Qiagen) and SYBR green (ThermoFisher). They were calculated as relative using the double delta Ct in comparison to Allstarneg expression levels. Expression levels of different isoforms in HeLa cells were determined in the same way and the results were analyzed with the formula 2delta Ct (GAPDH-target).

### HPV16 binding to cells and ECM

For HPV16 binding to cells, HeLa cells were seeded in 96-well optical-well plates at a density of 4000 cells per well and subsequently infected 16 h post seeding. About 1-2,5 ng AF568-labeled HPV16 were added per well. Cells were fixed with 4% PFA 2h p.i. and the actin cytoskeleton was stained with Atto647N-phalloidin for binding to cells experiments. Image stacks covering the whole cell were acquired as above, with a spinning disc microscope (Zeiss Axio Observer Z1, equipped with a Yokogawa CSU22 spinning disc module; Visitron systems GmbH, or a Nikon Ti2 eclipse equipped with a Dragonfly 600 spinning disc module, Oxford Instruments) using a 40x objective. SUM/MAX projections were then carried out with Fiji (ImageJ distribution). For the quantification of binding to cells, an outline of the cell area was generated using CellProfiler and used as a mask on the virus image to detect virus particles bound to cells. The number of particles in this image was then quantified using the Fiji plug-in 3D object counter. The number of cells per image were counted manually and the ratio of virus particles per cell was calculated.

For binding to ECM, HaCaT cells treated with NaClO_3_ for 24h were seeded at confluency (4×10^4^) in 96-well optical-well plates to generate ECM. HaCaT cells were detached 48h later by treatment with 20mM EDTA and ECM was extensively washed with PBS. About 1-2,5 ng AF488-labeled HPV16 were added per well. Samples were fixed with 4% PFA 2h p.i., washed with PBS, and subsequently blocked with 3% BSA in PBS, and stained with primary antibody against laminin-332 (AbCam). After washing, samples were incubated with AF568-labelled secondary antibody (Invitrogen) and washed again. Single slice images were acquired with a spinning disc microscope with a 40x objective (as above). For ECM binding, virus particle numbers were detected as above, and related to the intensity of laminin-332 staining.

### Synthesis of N-sulfated heparin

Heparin pyridinium salt was prepared as described (*41*). Briefly, heparin sodium salt (3.0 g) was dissolved in Milli-Q water (120 mL) and acidified by passage through AG50W-X4 resin (50 g). The aqueous solution was titrated with pyridine (1 mL) to a pH > 7 (between 8 to 9), then lyophilized to yield the pyridinium salt of heparin.

Fully desulfated heparin was prepared as described(*41*): briefly, the pyridinium salt of heparin (1 g) was dissolved in a DMSO:MeOH solution (100 mL, 9:1) and stirred under inert atmosphere at 100°C for 12 hours. After 12 hours, the reaction solution was brought to room temperature, diluted with Milli-Q water (300 mL) and cooled on an ice bath. The aqueous solution was then titrated with NaOH to a pH = 7. The solution was then evaporated until about 50 mL of solvent remained. The remaining solution was transferred to dialysis tubing (3,000 MWCO) and dialyzed over 48 hours with several changes of Milli-Q water, followed by lyophilization to yield fully desulfated heparin.

N-sulfated heparin with about 90% N-sulfation was prepared as described(*42, 43*). Briefly, fully desulfated heparin product (0.5 g) was dissolved in Milli-Q water (20 mL) and treated with sodium carbonate (1.25 g) and sulfur-trioxide pyridine complex (1.25 g). The solution was stirred at 48 °C for 24 hours. After 24 hours, additional sodium carbonate (1.25 g) and sulfur-trioxide complex (1.25 g) were added, and the solution was stirred at 48 °C for another 24 hours. After 24 hours, the solution was dialyzed as described above and lyophilized to yield ∼90% *N*-sulfated heparin by NMR analysis.

### Buffer and proteins

All AFM SMFS experiments were performed in phosphate buffered saline (PBS) buffer at pH 7.4, which was prepared by dissolving 1 PBS tablet (Medicago AB, Uppsala, Sweden) in 1 L milli-Q water (Millipore integral system, Molsheim, France) and filtered with 0.2 µm filters (Sarstedt, Germany) before use. Lyophilized streptavidin (Sigma Aldrich, St. Louis, MO) was dissolved in milli-Q water at 5 mg/mL and stored at −80°C. Thawed aliquots were used within a few days.

### Biotinylation of heparin and its derivatives

All heparin and its derivatives used in this study are listed in Table S1 together with their biochemical characteristics. All heparins were dissolved in milli-Q water at the desired concentration and gently shaken overnight at 4°C. Stock solutions of all heparins were aliquoted and stored at −20°C. Thawed aliquots were used within a few weeks.

For biotinylation, heparin and three selectively desulfated heparins were biotinylated via oxime ligation at their reducing end as described in (*44*). Briefly, all the heparins at stock concentration of 4 mM were first resuspended in coupling buffer (200 mM acetic acid/sodium acetate buffer, pH 4.8), separately. The solution was then mixed with aniline (80 mM; sigma) followed by EZ-link alkoxyamine PEG-4-SS-biotin (40 mM; PEG = polyethylene glycol, Thermo Fisher Scientific). All 4 reagents were mixed in equal volumes. The reaction was allowed to continue for 48h at 37°C. To remove unreacted biotin and aniline, the mixture was extensively dialyzed in milli-Q water using a membrane with molecular weight cutoff of 3.5 kDa (Spectra/Por, Thermo Fisher Scientific) at 4°C. The final products were aliquoted and stored at −20°C. The final concentration of biotinylated heparins was determined using the cetylpyridinium chloride (CPC) turbidity test as described elsewhere(*45*) and biotin quantification assay kit. Briefly, 50 µL of CPC reagent (0.2% w/v of CPC and 133 mM of MgCl_2_; both from Sigma) was mixed with 50 µL of heparins solution in a 96-well cleared bottom plate (VWR). The solution was then carefully mixed to avoid bubbles and incubated at room temperature for 30 mins. The absorbance was measured at 405 nm using a spectrometer (Varioskan® Flash, Thermo Fisher Scientific). The amount of biotinylation of the final products was quantified using commercial biotin quantification assay kit (QuantTag Biotin Quantification Kit-Vector Laboratories, CA) using the supplier’s protocol. Taking average molecular weight of GAGs (Table S1) and the concentration of GAG sample measured by CPC assay, the ratio between biotins and GAGs was found to be 1.02, indicating one biotin per GAG chain.

### Lipid vesicles

1-palmitoyl-2-oleoyl-glycero-3-phosphocholine (POPC) and 18:1 Biotinyl Cap PE (DOPE-Cap-B) were purchased from Avanti Polar Lipids (Alabaster, AL, USA). Small unilamellar vesicles (SUVs) were prepared from a mixture of POPC:DOPE-Cap-B (90:10, molar ratio) in PBS at a stock centration of 2 mg/mL by extrusion through a 50 nm polycarbonate membrane at least 21 times, using a mini extruder equipped with 1 mL syringe (Avanti Polar Lipids; 610020 and 610017) as previously described (*28*). Stocks were stored at 4°C under N_2_ and used within a few weeks.

### AFM based single molecule force spectroscopy (SMFS) measurements: Anchoring HPV16 virions to the AFM tip

HPV16 PsV particles were anchored to AFM cantilevers via PEG linker in a three steps process as previously described (*46, 47*). Firstly, amine groups are formed on the cantilevers using 3-(aminopropyl)trimethoxysilane (APTMS; Sigma-Aldrich). In the second step, the APTMS-treated cantilevered were conjugated with a PEG linker. Finally, the coupling of viruses to the PEG linker was performed.

To achieve this, three to four UV/ozone (UV Ozone Cleaner-ProCleaner™ Plus, Bioforce, IA, USA) treated (10 min) silica nitrate cantilevers (MSCT; Bruker AFM Probes, USA) were amino-functionalized with APTMS in the gas-phase salinization process as described elsewhere (*48*). Briefly, a desiccator (4 L) was filled with N_2_ gas for 2 min to remove any air and moisture. Then 30 µL of APTMS and 10 µl of triethylamine (Sigma) were pipetted into two small plastic trays, for example the lids of 1 mL Eppendorf tubes, and placed in the desiccator, which was filled with N_2_ again for 2 min. The cantilevers set on a cleaned Teflon sheet were immediately placed next to these small trays, the desiccator was sealed off, and filled with N_2_ for further 3 min. After 2h, the APTMS and triethylamine trays were removed and the desiccator was sealed off again, filled with N_2_ for another 2 min. The APTMS-treated cantilevers were kept in this environment at least overnight or until use.

Next, amino-treated cantilevers were carefully immersed in NHS-PEG-Acetal MW 2k (Creative PEG, USA) solution (1 mg dissolved in 0.5 mL chloroform and mixed with 30 µL of triethylamine) in a custom-made Teflon block holder. The holder was covered with a glass coverslip to avoid the evaporation of chloroform, and the cantilevers were incubated for 2h. Subsequently, the cantilevers were carefully rinsed with chloroform (3 x 10 min), gently dried with N_2_, and either stored under N_2_ until use or functionalized immediately with HPV16 particles.

Finally, the coupling of the PEG-acetal linker with HPV16 or fcHPV16 PsV particles was performed by immersing the PEG-coated cantilevers (∼1-to-2) in (w/v) 1% citric acid (Sigma) in milli-Q water for 10 mins. After washing with milli-Q water (3 x 5 min), the cantilevers were N_2_ dried. The cantilever was then installed on the cantilever holder and 70 µL of virus solution (∼2 x10^9^ TU/mL) was carefully pipetted on it. 2 µL of freshly prepared sodium cyanoborohydride (NaCNBH_3_; Sigma) solution (13 mg of NaCNBH_3_ dissolved in 20 µL of 20 mM NaOH (Sigma) and 180 µL of milli-Q water) was then immediately added to the virus solution on the cantilever, mixed very carefully and left to incubate for 40 min. After 40 min incubation time, 5 µL of 1 M (pH 8.0) ethanolamine (Sigma) solution was carefully mixed to the virus solution containing the cantilevers, and incubated for further 10 min. The cantilevers were then carefully rinsed either with 5x 1 mL RT PBS and were immediately used in AFM experiments.

### Preparation of model surface with biotinylated GAGs

Microscope glass cover slides with diameter of 22 mm (VWR, round, No.2) were cleaned using 7x detergent (MP Biomedicals, CA) and milli-Q water with 1:6 (volume) ratio, close to boil for 2h, rinsed thoroughly and stored in milli-Q water. Before use, the slides were rinsed with milli-Q water, N_2_ dried, and treated in UV/ozone for 30 min.

Supported lipid bilayers (SLBs) were formed by spreading of SUVs composed of a POPC:DOPE-Cap-B (90:10 molar ratio) mixture at concentration of 100 µg/mL on a cleaned glass cover slip, which was fixed on a custom-built stainless steel AFM holder using a bi-component Twinsil glue (Picodent, Germany) for creating a well that could hold a volume of about 100 µL. All incubation steps were performed in still solution. The excess sample at each incubation step was removed by repeated addition of a 2-fold increase of PBS and removing of excessive solution until the concentration of the solute was approximately 200 ng/mL. The solution was homogenized at each dilution step with at least 2 aspiration and release cycles. After forming SLBs, streptavidin was added to the well at the final concentration of 50 µg/mL for 30 mins. After a rinsing step, the streptavidin-coated SLB was exposed to biotinylated heparins at the final concentration of 5 µg/mL for 30 mins. For b-HA, final concentration was increased to 7 µg/mL while keeping the same incubation time to have similar number of chains in bulk solution. These conditions allowed the binding probability between HPV16 virion and heparin to be less than 20% for mainly probing the interactions between a single virion and a single heparin chain (see supplementary text S4 for details).

### AFM data acquisition and analysis

AFM-based SMFS measurements were performed using a NanoWizard 4 XP bioscience system (Bruker/JPK, Berlin, Germany) in PBS buffer at ambient temperature. Force-separation curves were recorded with virus-functionalized cantilevers; MSCT, probes C and D with nominal spring constant of 10 pN/nm and 30 pN/nm, respectively. The spring constant for each cantilever was calculated based on the thermal noise method(*49*), and found to be within 10% of the nominal spring constant value provided by the supplier. An approach speed was fixed at 1 µm/s and a retract speed ranging from 0.5 µm/s to 8 µm/s, with a maximum applied load of 600 pN, surface dwell time of 0 ms, and a ramp size of 400 nm, were used to collect force separation curves unless otherwise stated. All experiments were repeated at least twice, with separately but identical prepared AFM tips and heparins-coated model surfaces. Force-separation curves were analyzed using the JPK data processing software. The retraction curves were fitted with the freely jointed chain (FJC) model for quantitative analysis of the stretching of an individual PEG chain and to extract virus and heparin bond rupture forces with both persistence length and contour length as adjustable parameters. To avoid the contribution from non-specific tip-sample interactions, retract curves with rupture events occurring at a distance >10 nm were further analyzed. Instantaneous loading rates (*r*) were calculated by multiplying the effective spring constant, i.e., the slope of the FJC fit close to the rupture point, with the retract speed. Force histograms were computed and fitted with Gaussian to extract mean rupture forces. The kinetic parameters (*k*_off_ and *x*_β_) were calculated from the dynamic force spectra (DFS) data, which relates the mean unbinding force (*F*) and instantaneous loading rate (*r*). For this, we experimentally probed the interactions at various loading rates by varying pulling velocities in our SMFS experiments and fitted the DFS data (*F* versus *r*) with the Bell-Evans model(*50–52*) using linear regression analysis. The model is mathematically represented as,

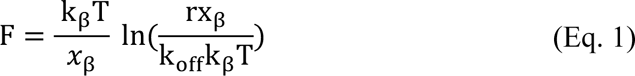

where *x*_β_ is the width of barrier and *k*_off_ is intrinsic transition or unbinding rate constant at F = 0. According to this model, a linear dependence of the mean rupture force as a logarithmic function of the instantaneous loading rate is predicted for stochastic bond rupture across a single energy barrier under external applied load(*50*). The standard error of mean rupture force was considered to determine the confidence interval for the kinetic parameters in this analysis.

To determine the rate of bond formation of our system or *k*_on_ analysis, we first measured the influence of contact time (the time the tip is in contact with the surface) on the binding probability (BP). BP is the measure of fraction of force curves displaying unbinding events. It is related with the interaction time (τ) with the following relationship:

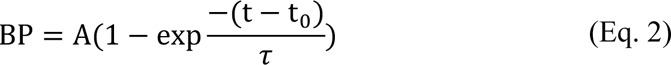

where A is the maximum measured BP and t_0_ is the lag time. BP was measured by collecting 500 force-distance curves for different contact (surface dwell) times of 0.05, 0.1, 0.15, 0.2, 0.3, 0.5, and 1s with keeping the rest of parameters fixed (extend/retract velocities = 1 µm/s, ramp size = 400 nm, maximum applied load = 600 pN). The plots of BP as a function of contact time were then fitted using the Origin software to extract τ using a least-squares fit of Eq. 2. Lag time was set to 2.18 ms for all data sets, which was determined from the fitting of BP plot of HPV16 and heparin data to Eq. 2 to set the same initial BP for all GAG conditions tested here. In the next step, *k*_on_was obtained using the following relationship between interaction time and effective concentration;

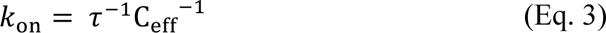

where effective concentration (C_eff_) represents number of binding molecules on an AFM tip within the effective volume (*V*_eff_) that are available for free equilibrium interactions. It is mathematically expressed as,

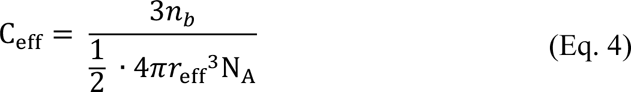

where *r*_eff_ is the radius of the sphere with volume, *V*_eff_ and is equal to equilibrium length of PEG linker ∼ 2 nm and the size of virion ∼50 nm (For details about how to calculate *r*_eff_, see(*53, 54*)), *n*_b_ is the number of potential bonds formed within half of sphere volume in which interaction can take place (this is set to 1 in our case because most of force-distance curve show only a single unbinding event) and N_A_, is the Avogadro’s number.

### Quantitation and statistical analysis Quantitation of HPV16 infectivity

Quantitation of HPV16 infected HeLa and HaCaT cells was performed by scoring GFP positive cells 48 hours p.i. by microscopy as described above for at least three independent experiments. Data was plotted with GraphPad Prism 7. Statistical analysis was performed using a Student’s t-test.

## Acknowledgments

We thank the members of the Schelhaas laboratory for critical comments on the manuscript. We acknowledge the facilities and technical assistance of the Umeå Plant Science Centre (UPSC) Microscopy Facility for access to AFM. We also thank the Biochemical Imaging Center at Umeå University and the National Microscopy Infrastructure, NMI (VR-RFI 2019-00217) for providing access to AFM and QCM-D. The funders had no role in study design, data collection, decision to publish or preparation of the manuscript.

## Funding

This study was supported by the German Research Foundation (DFG, grants SCHE 1552/6-1, SCHE 1552/3-2, INST 211/1029-1 to M.S.), the Knut and Alice Wallenberg foundation, the Swedish Research Council (VR grants 2017-04029; 2020-06242) and the Kempe foundations.

## Author contributions

Conceptualization: M.B. and M.S. Methodology: F.B, L.S.-M., M.B, and M.S. Investigation: F.B, L.S.-M., D. v. B. I.F.., K.T., A.B., and N.S. Visualization: M.S. Supervision: M.B. and M.S. Writing—original draft: F.B., L.S.-M., M.B, and M.S. Writing—review & editing: F.B., L.S.-M., M.B, N.S., and M.S.

## Competing interests

The authors declare no competing interests.

## Data and materials availability

All data are available on request.

## Notes

### Competing Interest Statement

The authors have declared no competing interest.

