## Supplementary material for "Site-specific sulfations regulate the physicochemical properties of papillomavirus-heparan sulfate interactions for entry": Suppl. Material

Fouzia Bano *et al.*

**This PDF file includes:**

Supplementary Text: S1 to S5
Figs. S1 to S7
Tables S1 to S3
References (1 to 5)

### Supplementary Text

#### S1. Perturbing expression levels of HS-biosynthetic enzymes in cells

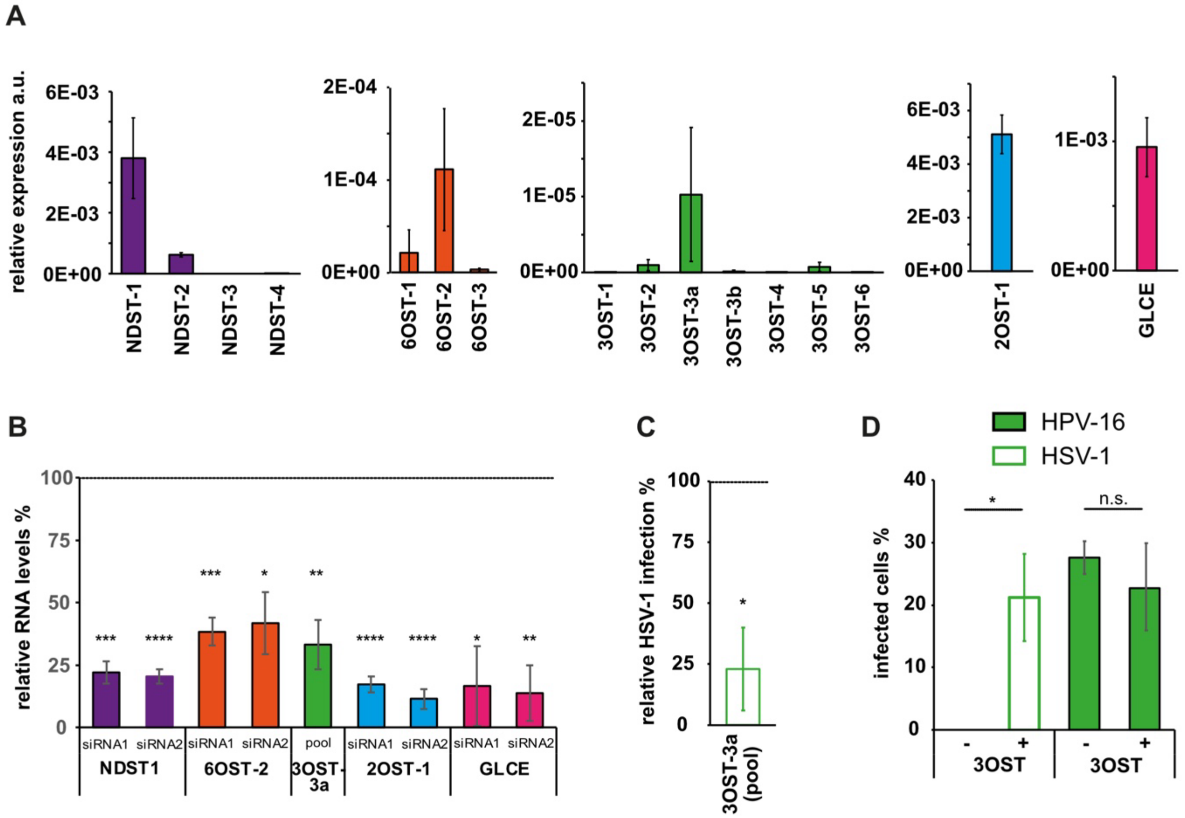


**Fig. S1**. **Perturbing expression levels of HS-biosynthetic enzymes in cells.**

(**A**) mRNA levels of all enzymes involved in the biosynthesis of sulfation pattern of HS were assessed via quantitative PCR (qPCR). mRNA levels for indicated enzymes of least three independent passages of HeLa cells were normalized to a housekeeping gene (GAPDH) and depicted as arbitrary units (a.u.) ± SD. (**B**) mRNA levels of HS modifying enzymes after RNAi (as in Fig. 1) were assessed by qPCR. The mRNA levels of these isoforms in transfected cells are depicted relative to non-target control (line). Shown are averages of three independent experiments ± SD. (**C**) HeLa cells were transfected with siRNAs against 3OST-3a (15 nM, pool). Two days post transfection (p.t.), cells were infected with HSV-1-GFP and fixed 6h p.i.. Shown are average infection levels relative to control siRNA (15nM, line, 100%) ± SD. (**D**) CHO cells were transfected with expression constructs of 3OST-3a-eGFP or eGFP (control) and infected with HSV-1 or HPV16-PsV. Shown are the amounts of infected cells (in %) expressing 3OST-3a or GFP of three independent experiments ± SD. For all quantifications, a two-tailed student's t-test was performed with $p < 0.05$ (*), 0.01 (**), 0.005 (***), 0.0001 (****) or non significant (n.s.).

**S2. Heparin derivatives used in this study.**

**Table S1**: Heparins used in this study.

| **Name** | **Molecular weight (kDa)** | $n_{ds}$§ | **Sulfate group per disaccharide#** |
|---|---|---|---|
| **Heparin** | 15 | 25 | 2.42 |
| **2O-desulfated heparin (2O-deS)** | 15 | 25 | 1.75 |
| **6O-desulfated heparin (6O-deS)** | 15 | 25 | 1.48 |
| **N-desulfated heparin (N-deS)** | 15 | 25 | 1.55 |
| **b-Hyaluronan (b-HA)** | 23 | 57 | 0 |

§Values for number of disaccharide unit per chain estimated from the average molecular weight of heparin.

#Average number of sulfate groups per disaccharide calculated from disaccharides analysis provided by the supplier.

### S3. HPV 16 binding to the extracellular matrix

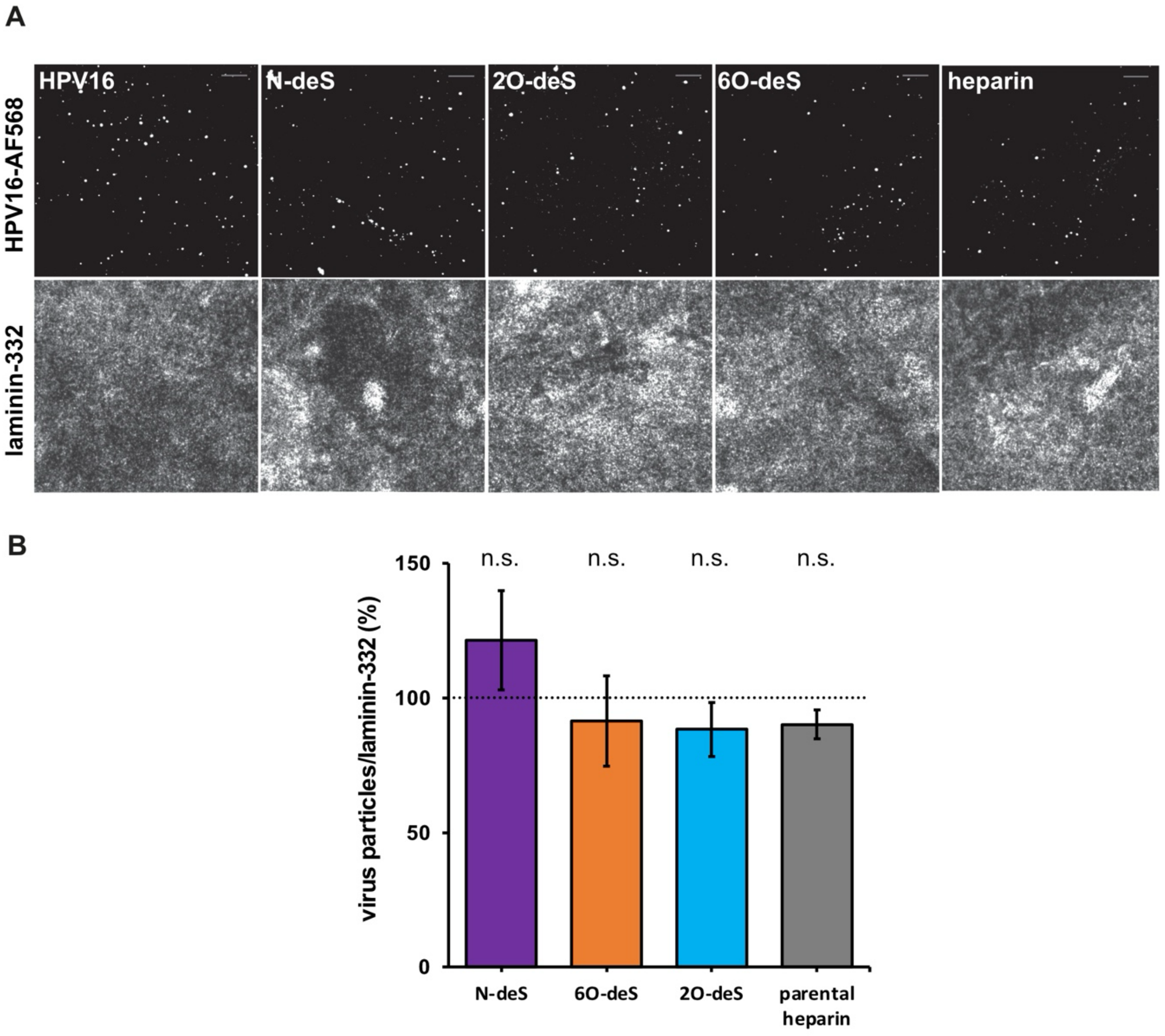


**Fig. S2**. **HPV16 binding to ECM after incubation with heparins.**

(**A**) fluorescently-labeled HPV16 (AF-568) were pre-incubated with the indicated glycans for 1h at 1mg/ml concentration and subsequently allowed to bind to undersulfated ECM (here indicated by laminin-332 staining) for 1h prior to fixation. (**B**) Samples were imaged by confocal microscopy and analyzed using Fiji. The number of particles were normalized to the amount of laminin-332 and are depicted relative to the control, i.e., heparin. Scale bar is 10 µm. For the quantification, a two-tailed student's t-test was performed with $p < 0.05$ (*), 0.01 (**), 0.005 (***), 0.0001 (****) or non significant (n.s.).

**S4. Characterization of the biomimetic platform by quartz crystal microbalance with dissipation (QCM-D) to study HPV16 binding to heparin**

***QCM-D experiments***

QCM-D sensors with fundamental frequency of 5 kHz and silica coating (AWS SNS 000049A) were purchased from Advanced Wave Sensors (AWS) S.L., Spain. The sensors were cleaned with 2% (w/v) sodium dodecyl sulfate (SDS) for 1h, rinsed thoroughly with milli-Q water, blown dry with N2, and treated in a UV/ozone cleaner (Bioforce Nanoscience, Ames, IA) for 30 min before use.

QCM-D measurements were performed with an X4 system (AWS, Spain) equipped with four independent channels. The system was operated with a flow rate of 20 µL/min unless otherwise stated using software-controlled fluidic modules at working temperature of 22ºC. The QCM-D response (frequency shift Δf and dissipation ΔD) was recorded at six overtones ($i$ = 3, 5, 7, 9, 11, 13) and data for $f_i$ = 3 is presented, unless otherwise stated. All experiments were performed in phosphate buffered saline (PBS) buffer at pH 7.4, which was prepared by dissolving 1 PBS tablet (Medicago AB, Uppsala, Sweden) in 1 L milli-Q water (Millipore integral system, Molsheim, France) and filtered with 0.2 µm filters (Sarstedt, Germany) before use. The buffer was degassed using a sonicator (Elmasonic, S40H, Singen, Germany).

***Platform characterization***

QCM-D simultaneously measures the shift in resonance frequency (Δf), and dissipation (ΔD) upon exposing a sensor surface to molecules of interest. Both Δf and ΔD are sensitive to "wet" mass density of molecular film which includes dry mass of surface bound molecules with hydrodynamic-coupled water and the mechanical properties of the film(*1*).

To ensure similar properties of the heparin layers constructed as described in Supplementary Fig. 3A, mass adsorption in real time and mechanical properties of the grafted heparin film were measured and delineated using QCM-D (fig. S3B). These measurements demonstrated biotin-specific formation of stable heparin films (fig. S3B and Table S3). Similar profiles of parametric plots of ΔD/–Δf ratio (a relative measure for film softness) as a function of –Δf (a relative measure for film surface density) suggested that for a given surface density, all heparin films have comparable softness (fig. S3D). Moreover, QCM-D measurements confirmed that HPV16 specifically interacted with the heparin film (fig. S3B), as no binding was observed on hyaluronic acid (HA) films, an unsulfated GAG, used as a negative control (fig. S3C). Thus, all heparin films were stably immobilized and mechanically comparable, and, importantly, allowed specific GAG-HPV16 interactions.

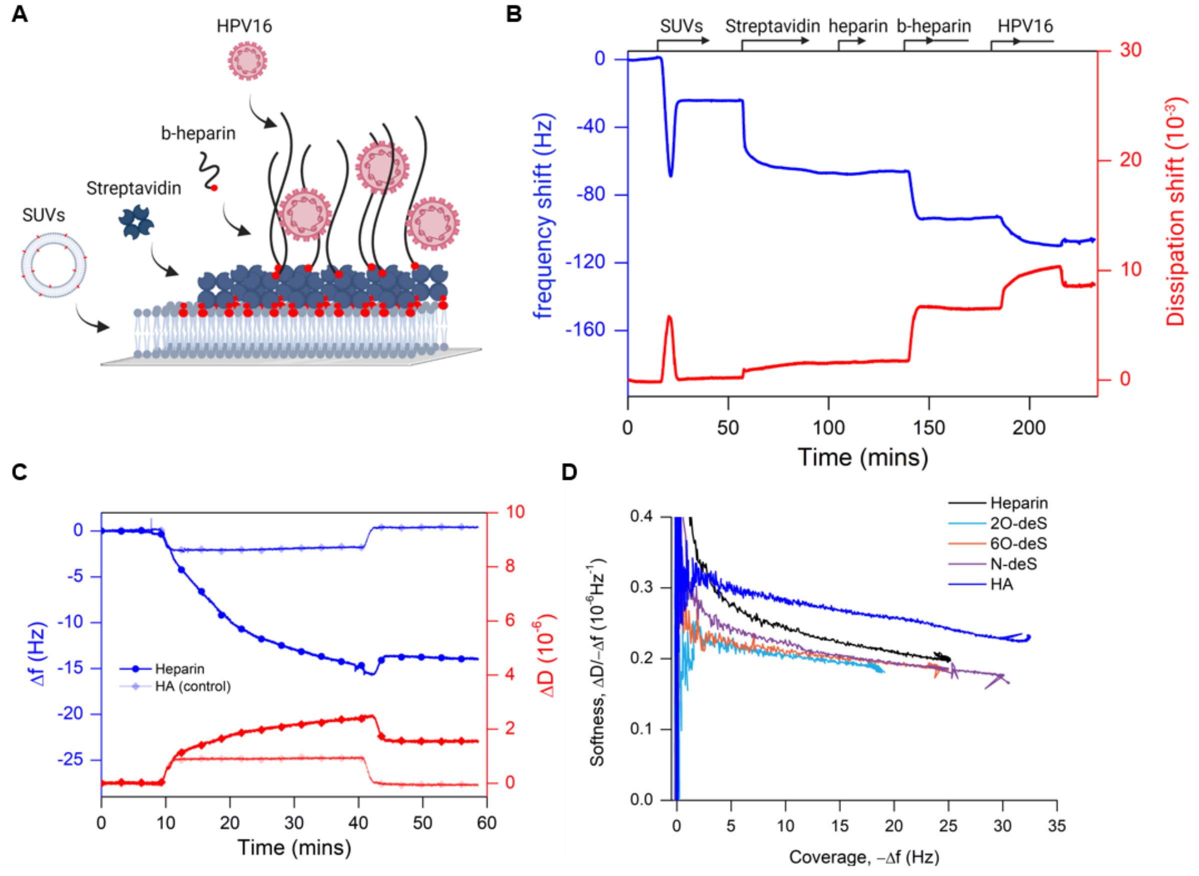


**Fig. S3**: **Heparin film formation and HPV16 binding observed by QCM-D**.
(**A**) Illustration of the design and step-by-step assembly of a surface displaying end-grafting immobilization of heparin via biotin on a monolayer of streptavidin formed on a supported lipid bilayer (not to scale). The supported lipid bilayer is formed by substrate-induced fusion of small unilamellar vesicles (SUV). (**B**) Representative QCM-D plot displaying shift in frequency (blue) and dissipation (red) for the formation of a stable HPV16-binding heparin film. Arrows on top of the graph indicate a continuous flow of biomolecules. A line following an arrow indicates continuous flow for 20 minutes followed by an additional 10 min incubation without flow. Empty time slots represent rinsing with PBS buffer under flow. (**C**) QCM-D plots comparing the

binding of HPV16 to heparin and HA (control surface). (**D**) Parametric plot for binding data of the four biotinylated-heparin variants and biotinylated-HA to streptavidin as indicated. Curves displaying softness (ΔD/–Δf) as a function of coverage, represented as frequency shift, –Δf for heparin, its derivatives, and HA had comparable shapes. The softness decreased with an average slope of $-2.6 \times 10^{-3}$ for all films as a function of surface coverage, implying that overall, all films display a comparable softness at a given surface coverage.

**Table S2**: QCM-D frequency and dissipation shifts for biotinylated films of heparin and its derivatives on streptavidin. Average values ± S.D. were obtained from 2 independent experiments.

| GAGs | Frequency shift, Δ*f* (Hz) | Dissipation shift, Δ*D* ($10^{-6}$) |
|---|---|---|
| Heparin | -22.1 ± 3.4 | 4.2 ± 0.6 |
| 2O-deS | -20.7 ± 2.5 | 3.5 ± 0.3 |
| 6O-deS | -25.9 ± 3.3 | 4.6 ± 0.4 |
| N-deS | -25.9 ± 2.0 | 4.4 ± 0.2 |
| HA | -36.8 ± 0.5 | 8.2 ± 0.05 |

**S4. Single-molecule nature of unbinding events and specific nature of HPV16-heparin interactions by AFM-based SMFS**

Binding events between am AFM tip carrying a single virion immobilized via a polyethylene glycol (PEG) linker (fig. S4) recorded by AFM-based SMSF were deemed to correspond to a specific interaction between a single virion and a single heparin chain because: (i) the binding probability (BP) of specific unbinding events was between 4 and 11%, with multiple unbinding events observed in <2% of curves (fig. S5B, filled black bars); a criteria used for predicting stochastic binding interactions at the single-molecule level(*2, 3*), (ii) fitting of a single rupture event of FD curves with the freely jointed chain (FJC) model revealed an average and velocity-independent Kuhn length of 7.9 ± 3.1 Å, which is in agreement with the literature of 7 Å for the stretching of a single PEG chain in PBS(*4*) (fig. S5C), and (iii) the PEG stretching parts prior to rupture event in every case fall onto a single master curve upon normalization according to predictions of the polymer stretching model (fig. S6); a measure to identify rupture events coming from the stretching of the same type of molecule, which is a single PEG chain in our case. Furthermore, the binding specificity between HPV16 and heparin was verified by (i) the introduction of free heparin as a competitor (fig. S5B, empty cyan bars), (ii) a tip lacking HPV16 (i.e., PEG linker only, fig. S5B, empty grey bars, fig. S5D), and (iii) a HA surface (fig. S5B, empty blue bars).

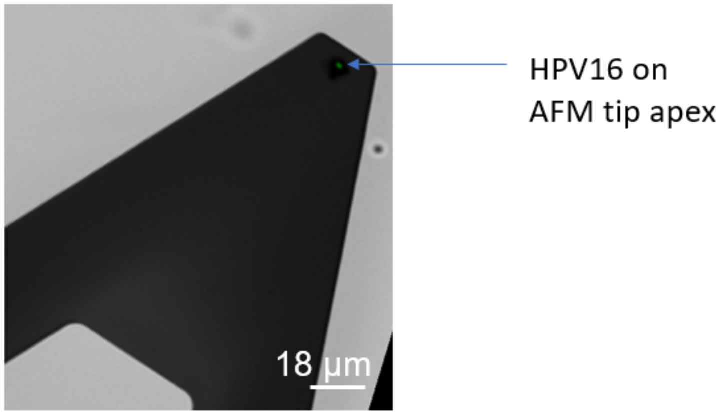


**Fig. S4**. **A single HPV16 virion on an AFM tip.**
Composite brightfield and a fluorescent image showing a single fluorescently labelled HPV16 particle on an AFM tip apex.

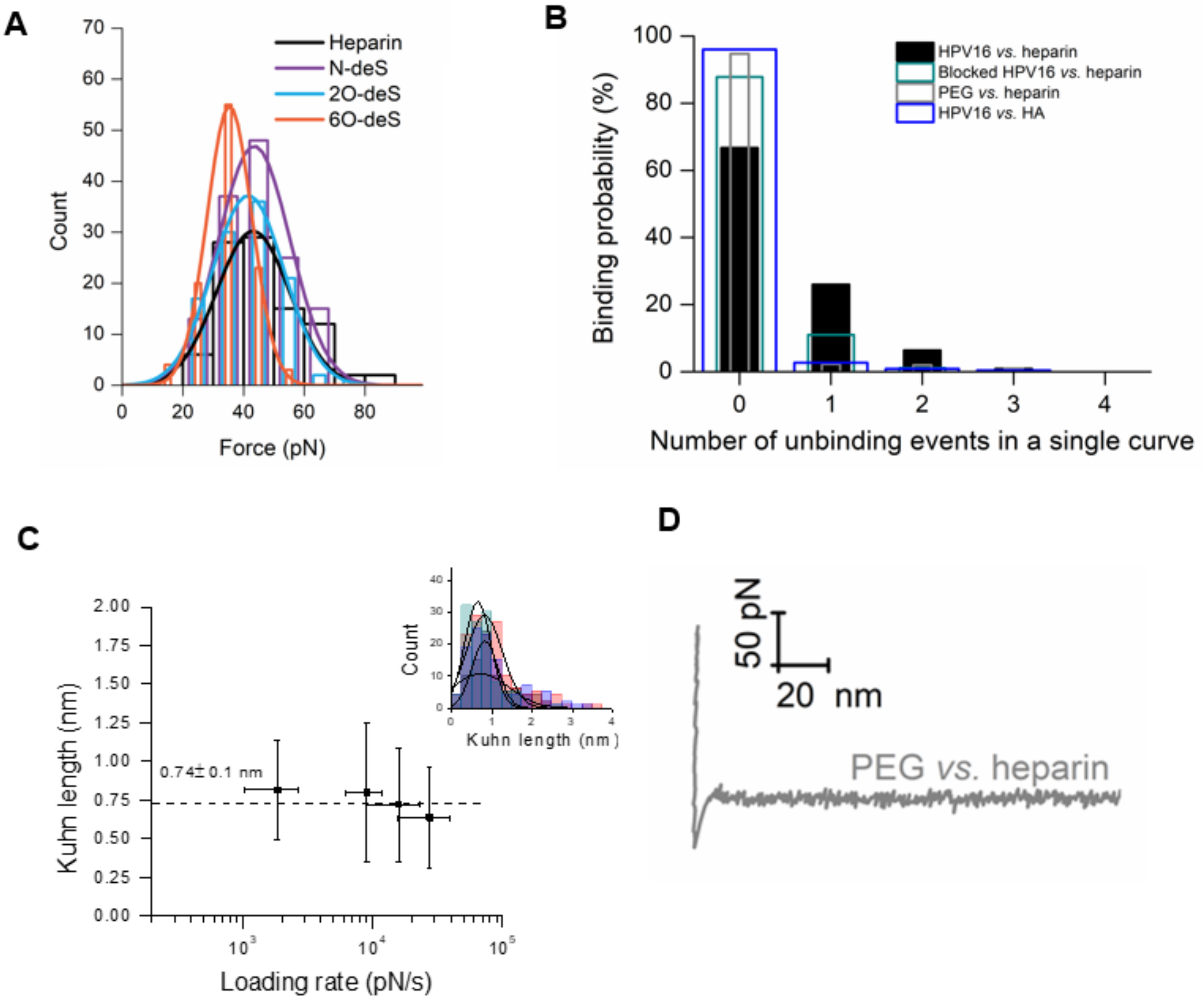


**Fig. S5. Supplementary force data for the specificity and confirmation of the single unbinding events.**

(**A**) Force histograms with Gaussian fits to rupture force histograms for loading rate of 8046 ± 3380 pN/s (heparin), 8864 ± 3897 pN/s (N-deS), 11080 ± 3891 pN/s (2O-deS) and 8479 ± 4603 pN/s (6O-deS) for the binding interactions between HPV16 and indicated heparin surfaces. (**B**) Binding probability for the conditions as indicated. For the blocking assay, free heparin (15 kDa) was introduced to the chamber at 20 µg/mL. A total of 1000 curves at retract velocity of 1 µm/s were collected for each condition. (**C**) Kuhn length versus instantaneous loading rates. Dash line represents the mean value ± S.D. the inset shows the histograms of the Kuhn length for four selected loading rates with Gaussian fits that were used to calculate the mean and standard deviations. (**D**) Selected retract curve registered at 1 µm/s for probing the interactions between a virus-lacking tip (PEG-coated) and a heparin surface, showing that no specific event is observed at >10nm from the contact point.

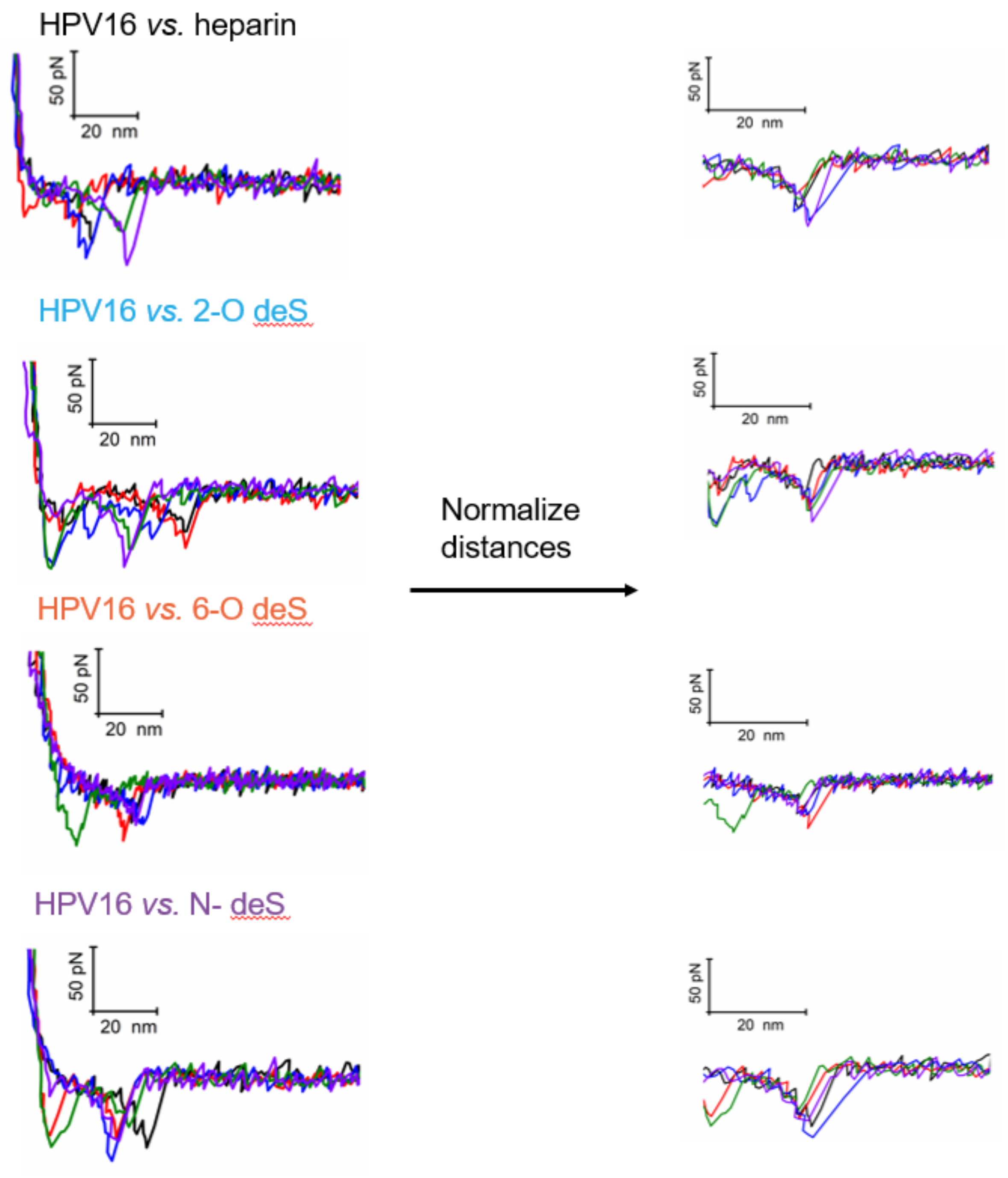


**Fig. S6. Normalization of force curves to further confirm single unbinding events.**
Five randomly selected force-distance curves recorded at 1 μm/s for four heparin surfaces and conditions as stated in Fig. 6b displaying a single unbinding event at various distances alongside with non-specific events at very small distances. Displayed on the right side are the same curves normalized such that $x_{norm} = {}^{x}/_{x^*}$ with $F(x^*) = F^*$, where $F^*$ was set for each heparin close to rupture events. According to

a polymer stretching model (FJC in our case), if all the normalized curves overlapped, then the pulling of a single PEG chain has been probed(*5*). This is indeed the case for our system.

### S5. Force histograms overview for HPV16-heparin interactions

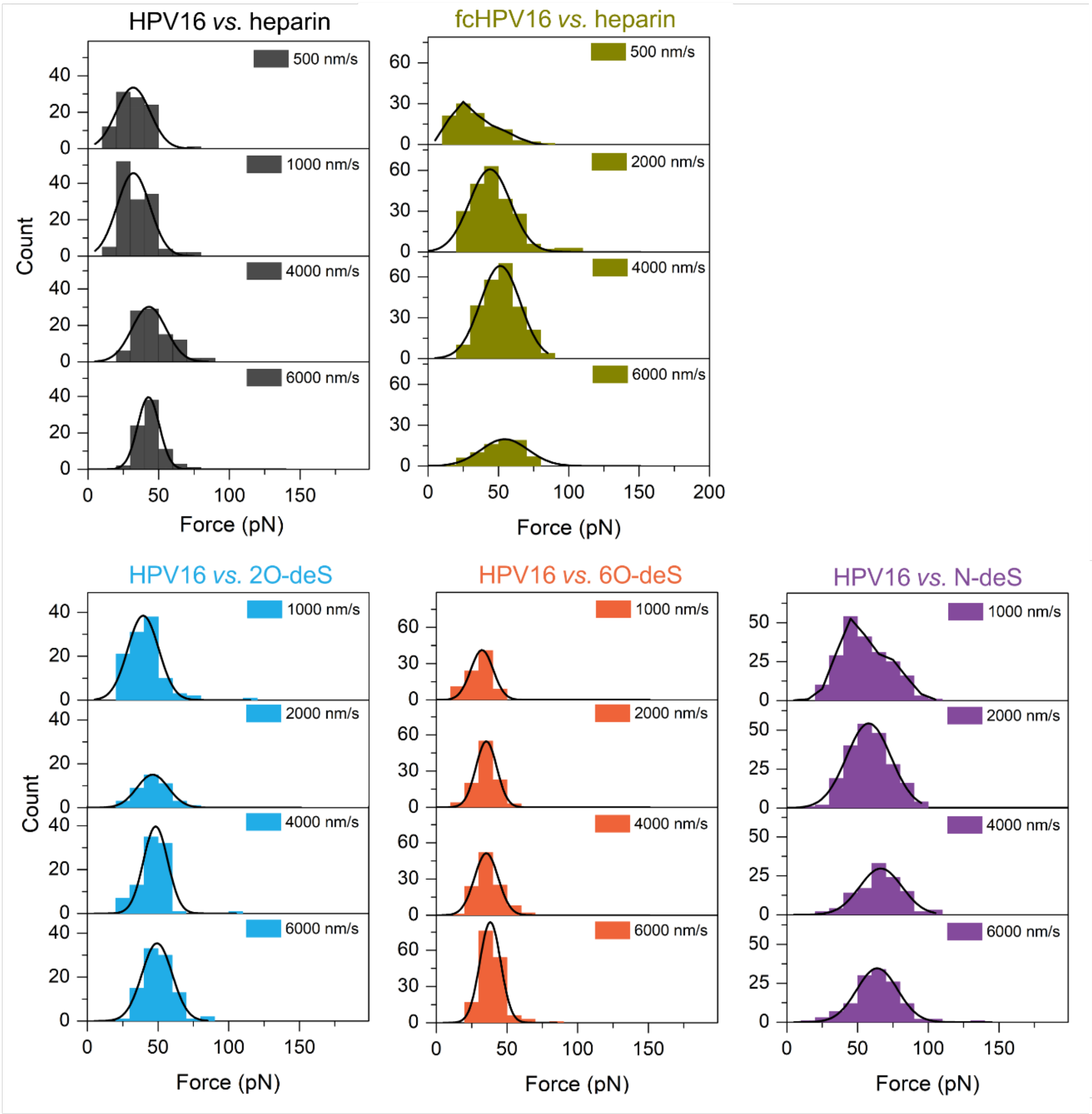


**Fig. S7. Histograms of rupture (unbinding) forces for HPV16 and heparin interactions.**

Force histograms for the interactions between HPV16, fcHPv16 and indicated heparin surfaces registered at different retract velocities (~1000 curves for HPV16 and heparin and ~3000 curves for the rest of conditions).

**Table S3:** Summary of kinetic parameters for the interactions between HPV16 and four heparin surfaces obtained from fitting of dynamic force spectra data with Bell-Evans model and binding probability analysis. N. D. = not determined

| PsV on AFM tip | Heparin surfaces | $k_{\mathrm{off}}$ ($s^{-1}$) | $x_{\beta}$ (nm) | $k_{\mathrm{on}}$ ($10^6\ M^{-1}s^{-1}$) | $K_{\mathrm{D}}$ (nM) |
|---|---|---|---|---|---|
| **HPV16** | Heparin | 0.004 ± 0.003 | 1.22 ± 0.38 | 2.03 ± 0.63 | 1.96 ± 2.03 |
| | 2O-deS | 0.015 ± 0.004 | 1.18 ± 0.07 | 3.20 ± 0.79 | 4.69 ± 1.17 |
| | 6O-deS | 0.12 ± 0.04 | 1.19 ± 0.06 | 4.61 ± 1.37 | 26.1 ± 12.6 |
| | N-deS | 12.2 ± 5.2 | 0.40 ± 0.06 | 1.43 ± 0.26 | 8536 ± 3950 |
| **fcHPV16** | Heparin | 18.7 ± 6.6 | 0.36 ± 0.01 | N.D. | N.D. |
